# Intracellular and extracellular dynamics of herpes simplex virus 1 DNA and infectious particles in epithelial and neuronal cells

**DOI:** 10.1101/2025.05.13.653775

**Authors:** Shadisadat Esmaeili, David A. Swan, Keith R. Jerome, Joshua T. Schiffer, Marius Walter

## Abstract

Herpes simplex virus 1 (HSV-1) infection of epithelial cells is lytic, while infection of neurons typically results in long-term latency. However, the rates at which HSV-1 replicates and spreads in epithelial cells versus neurons under low and high multiplicity of infection (MOI) conditions remain undefined. Identifying these rates requires the application of mathematical models to carefully designed viral kinetic experiments. It is also critical to differentiate the dynamics of infectious viral particles versus viral DNA, as both quantities are routinely measured in *in vitro* experiments and human studies using plaque assays and polymerase chain reactions, respectively. Here, we developed mechanistic mathematical models to describe HSV-1 dynamics after infection of epithelial Vero cells and neuronal N2A cells, at high (3) and low (0.01) MOI. Our model recapitulates the dynamics of cell-free and cell-associated viral DNA and plaque-forming units (PFU). In epithelial cells, the model describes a pre-productive eclipse phase with a mean duration of 10.9 and 12.8 hours prior to HSV DNA replication and PFU production, respectively. Cells exited the eclipse phase as early and late as 2.5 and 32 hours, respectively. Infected cells produced a single PFU for every 224 HSV DNA genomes. PFU egressed at a constant rate, whereas the HSV DNA egress rate increased over time, before saturating at a 15 times higher rate. Under low relative to high MOI conditions, Vero cells spent 7 hours longer in the eclipse phase, had a 12-hour delay prior to egress, and had a longer mean duration of productive infection (14 versus 3.5-hour half-life). Secondary epithelial cell infection in low MOI experiments was overwhelmingly due to cell-to-cell viral spread and originated from a small number of early-producer cells. Neuronal cells produced viruses at a 5-fold lower rate and had a longer (mean: 42 hours) and more variable eclipse phase, with some neurons remaining in eclipse for more than a week. Our results highlighted large differences in HSV egress rates, as well as infected cell eclipse phase duration and death rates, in epithelial cells versus neurons during low and high MOI infection. The observed viral dynamics in neurons reflect a balance between active replication and latency.

## Introduction

Herpes simplex viruses (HSV) 1 & 2 are persistent double-stranded DNA viruses that infect close to 70% and 15% of the human population, respectively^1^. HSV infection causes oral and genital ulceration that can have severe physical and psychosocial consequences, but lacks vaccines and eradication strategies. During primary infection, HSV replicates within and spreads between epithelial cells, leading to skin or mucosal erosions. HSV travels to ganglionic neuronal cell bodies via peripheral nerve endings, where it remains latent for life^2^. Periodic reactivation of HSV in the ganglia results in the transport of infectious virus back to the skin and mucosal sites, leading to ulcer formation and frequent asymptomatic shedding^3–5^. HSV is lytic to epithelial cells but establishes a lifelong latent infection in infected neurons. A detailed quantitative assessment of replication kinetics in these two cell types has not yet been performed.

Viral dynamic models are a valuable tool for deciphering the mechanisms underlying observed viral load trajectories. These mathematical models typically consist of deterministic or stochastic differential equations applied to viral load data from in vitro experiments^6–10^, animal models of infection^11,12^, or human cohorts^5,13–17^. The purpose of viral dynamic models is to couple non-linear interactions between replicating viruses, target cell depletion, immune responses, and therapies to derive mechanistic conclusions, which can sometimes be used to improve therapies^14,18,19^. For HSV-2, models applied to human shedding and immunologic data demonstrated that viral release from neurons is nearly continuous^4^; density of tissue-resident T cells predicts HSV-2 viral load in tissue microenvironments^5^; viral rebound is due to seeding of new mucosal micro-environments^17^; rapid spread of antiviral cytokines from activated T cells rather than direct TCR-mediated killing leads to elimination of most HSV-2 infected cells^16^; overall immune status predicts HSV-2 shedding rates^13^ in people living with HIV; T cell imprints in tissue are a manifestation of prior shedding rather than future protection^20^; HSV-2 reactivations are not predictable in individuals^15^ based on past shedding; and antivirals must have a long half-life to significantly lower HSV-2 shedding rate^14,21^.

A limitation of most existing viral dynamic models is that they vastly oversimplify the viral replication process due to the limitations of available data. Only a few existing models differentiate rates of production and clearance of viral genomes measured with polymerase chain reaction (PCR) from infectious viral particles measured by plaque assay^22^. Most models fail to discriminate the kinetics of cell-associated and cell-free viral particles, overlook the potential high importance of initial viral conditions, assume exponentially distributed lifespans for each cellular stage of infection without assessing other possible distributions, and do not contrast infection in cell types which favor lytic versus latent infection. In addition, high and low multiplicity of infection (MOI) conditions may lead to dramatically different infection cycles within individual cells, which is also typically overlooked.

An accurate description of the viral replication cycle requires more detailed mathematical models, but also highly granular experimental data designed explicitly for these models. Here, we record the dynamics of infection following high and low MOI infection of epithelial and neuronal cell lines. We quantify intra- and extracellular levels of HSV-1 DNA and infectious particles over time and use this rich experimental dataset to compare dozens of mathematical models. Our results demonstrate much higher rates of production and cellular egress for viral DNA relative to infectious particles. We observe that high MOI conditions favor a shorter pre-productive eclipse phase, shorter time to viral egress, and lower infected cell lifespan relative to low MOI conditions in epithelial cells. We find that cell-to-cell spread is the predominant mechanism of cellular infection in epithelial cells. Finally, we highlight that epithelial cells have much shorter and less variable pre-productive eclipse phases compared to neurons. This likely reflects separate outcomes between the two cell types following infection, one leading exclusively toward lytic replication and the other towards a balance of lytic infection and long-term latency.

## Results

### Cell-associated and cell-free viral DNA and infectious particle kinetics in Vero and N2A Cells following low and high MOI HSV-1 infection

We modeled the dynamics of HSV-1 in cell culture. Models were fitted to experimental data after infection of African green monkey epithelial Vero cells and murine neuroblastoma N2A cells. In Vero cells, 3 × 10^5^ cells were infected with HSV-1 at MOI=3 (high MOI) and MOI=0.01 (low MOI), in three biological replicates. Cell-free and cell-associated viral DNA were measured over time by quantitative polymerase chain reaction (qPCR), after collection of the supernatant and cell pellet, respectively. In addition, in the high MOI experiment, the titer of infectious particles, or plaque-forming units (PFU), in the supernatant and cells was measured by plaque assay (**Fig 1A, B**). While plaque assay only detects infectious viral particles, qPCR detects infectious and non-infectious HSV DNA, as well as DNA replication intermediates and other DNA fragments generated during the replication process or released after cell death^23^.

**Figure 1.**
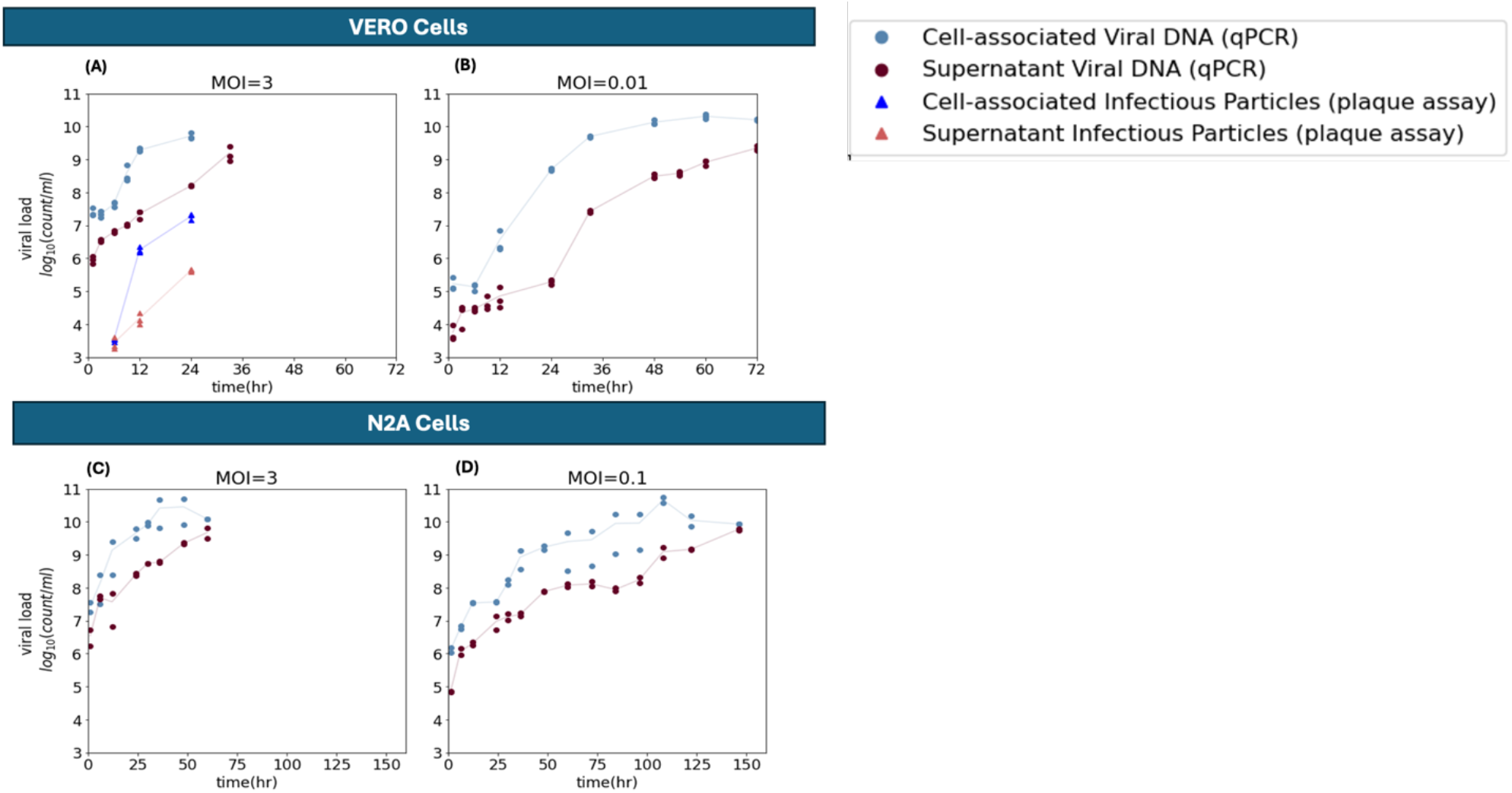
Viral dynamics in Vero cells and N2A cells. A, B: Dynamics of cell-associated and cell-free viral titers, as measured by qPCR and plaque assay after infection of Vero cells at (A) high (MOI = 3, left) and (B) low MOI (MOI = 0.01, right). n=3 biological replicates at each time point. C, D: Dynamics of cell-associated and cell-free viral DNA titers, as measured by qPCR after infection of N2A cells at (C) high (MOI = 3, left) and (D) low MOI (MOI = 0.1, right). n=2 biological replicates at each time point. qPCR data is expressed in copies per mL, and plaque assay data is expressed in plaque-forming units per mL (PFU/mL). Note differences in the x-axis time scale in panels A & B vs C & D, reflecting slower viral kinetics in N2A cells.

In Vero cells, viral DNA levels measured by qPCR were about three orders of magnitude higher than PFU titer of infectious particles measured by plaque assay **(Fig 1A)**. In addition, viral DNA (**Fig 1A, B**) and infectious titers (**Fig 1A**) were one to two orders of magnitude higher in cells than in the supernatant, indicating that HSV-1 remains principally cell-associated during lytic infection. In the high MOI experiment (MOI=3), almost all cells were infected at the beginning and the infection progressed rapidly, until maximum cytopathic effect after 24h (**Fig 1A**). In the low MOI experiment (MOI=0.01), the infection started in a small fraction of cells and progressed for 72 hours, until all remaining target cells were infected. Cell-associated viral DNA levels began to plateau after 36 hours (**Fig 1B**), compared to 12 hours following infection at MOI=3 (**Fig 1A**). Cell-free viral DNA levels remained low for the first 24 hours, followed by a period of rapid increase between 24 and 48 hours, and finally reached maximum levels after 72 hours (**Fig 1B**).

These experiments were repeated in mouse neuronal N2A cells (**Fig 1C, D**). 10^6^ cells were infected with HSV-1 at MOI=3 (**Fig 1C**) and 0.1 (**Fig 1D**), in two biological replicates. N2A cells required higher inoculation levels to initiate productive infection and an MOI of 0.1 was used for the low MOI condition (**Fig 1D)**. Cell-free and cell-associated viral DNA levels were measured over time by qPCR. Like in Vero cells, cell-associated levels were around 1.5-2 orders of magnitude higher than titers in the supernatant (**Fig 1C, D**). The infection progressed at a significantly slower rate in N2A cells than in Vero cells. At MOI =3, the infection lasted for 60 hours until all cells became infected and started dying, compared to 24 hours in Vero cells.

Furthermore, even with a higher MOI (0.1 in N2A vs 0.01 in Vero), the viral load increased until 150 hours in N2A cells compared to 72 hours in the Vero cells. This showed that HSV-1 infection progressed more rapidly in epithelial Vero cells versus neuronal N2A cells.

### Optimal model selection based on fit to HSV-1 kinetic data following Vero cell infection

By distinguishing infectious viral particles from viral DNA, and cell-free from cell-associated viruses, our experimental system offered a rich template for modeling viral dynamics. Mathematical models usually consider only a single population of viruses. Here, we developed multiple competing models to simultaneously recapitulate the dynamics of four viral populations, namely cell-free and cell-associated infectious particles, and cell-free and cell-associated viral DNA. We employed a three-stage approach to rule out competing models and identified an optimal model that closely reproduced the viral kinetic data of **Fig 1**. This approach is summarized below and presented in **Fig 2**.

**Figure 2.**
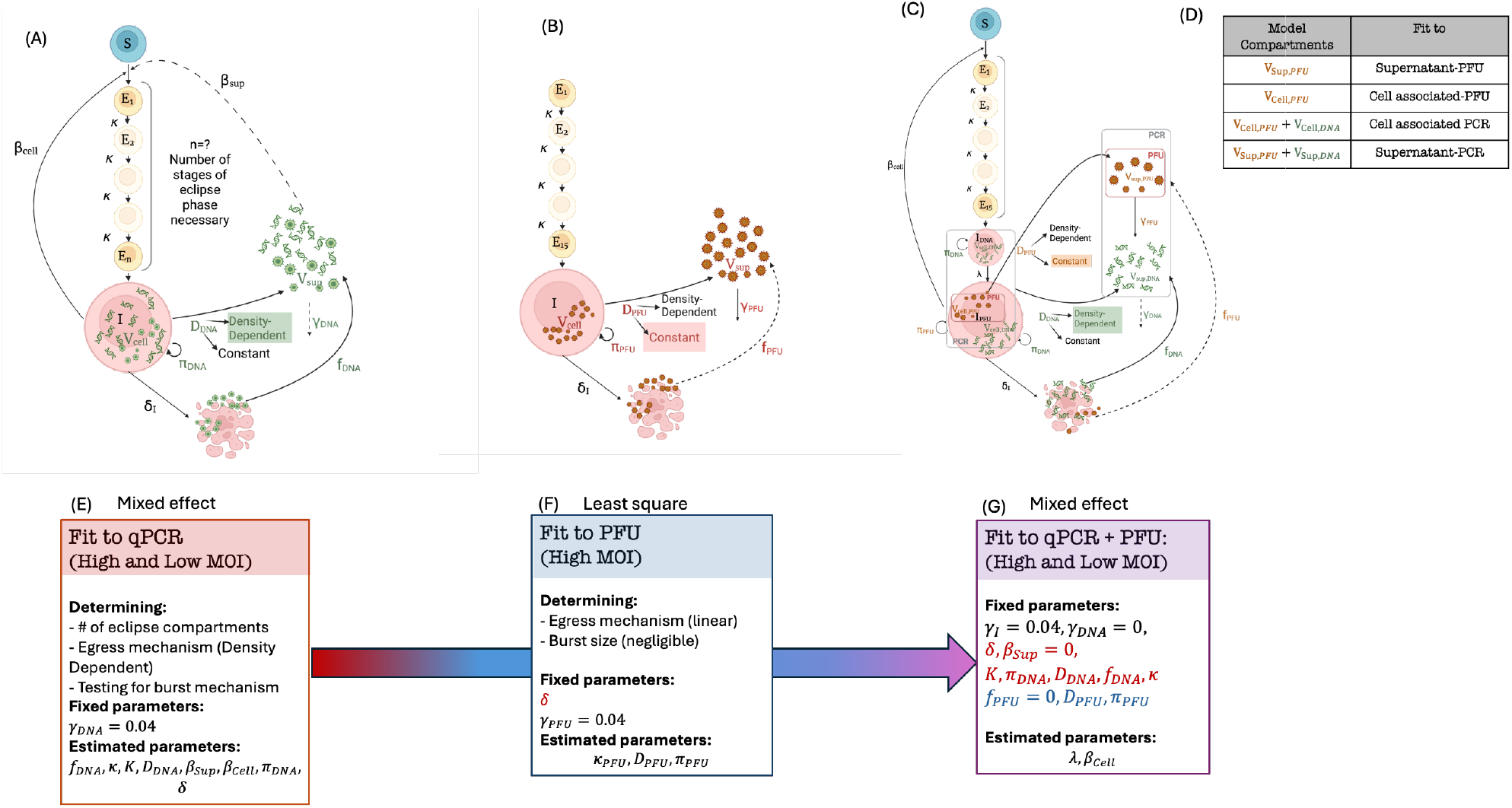
Optimized mathematical model schematic for Vero Cells and fitting approach: Schematic of the mathematical model fit to (A) high and low MOI qPCR data, (B) high MOI PFU data, (C) concurrent high and low MOI qPCR and PFU data with susceptible cells (*S*), cells in the eclipse phase (*E*_*i*=1,*…,n*_), and productively infected cells (*I*), which generate viral particles at rates *π*_*DNA*_(and *π*_*PFU*_). Viral particles egress to the supernatant at a constant or density-dependent diffusion rate. The susceptible cells are infected through cell-to-cell contact or by free virions in the supernatant at rates *β*_*Cell*_ and *β*_*Sup*_. Infected cells die at the rate *δ*_*I*_ and burst. Viral DNA and PFU inside the cell then enter the supernatant with burst-size *f*_*DNA*_ and *f*_*PFU*_, respectively. The dashed arrows indicate mechanisms that were initially considered in the model and then removed due to their negligible estimated parameter values. The susceptible compartment (S) was removed from the PFU model (B) since almost all cells are immediately infected at high MOI. In the final model, the production of infectious virions is delayed by the transition rate *λ* (D) Mapping model compartments to qPCR or plaque assay data shown in Fig 1. (E) Fitting the model to high and low MOI qPCR data using non-linear mixed effect methods to determine the shape of the distribution of the half-life of the infected cells in the eclipse phase as well as the mechanism of the viral DNA diffusion into the supernatant. All parameters associated with the dynamics of viral DNA were estimated while fixing the clearance rate of the free virions (*γ* = 0.04). The first data point (at t=1) was used as the initial viral load. (F) Fitting the model to plaque assay data and estimating all other parameters associated with the dynamics of PFU data except *γ. δ*_*I*_ was fixed to the value from the high MOI experiment estimated in (E). (G) Fitting the combined model for infectious and non-infectious viral particles to qPCR and plaque assay data. All DNA and PFU-related parameters were carried forward from stages (E) and (F), respectively. Only *λ* and *β*_*Cell*_ were estimated at this stage. (A-C Created in https://BioRender.com)

### Three-stage model development and sequential data fitting

The optimal model was identified using a three-stage approach. **Stage 1** fit viral DNA using qPCR data only, **Stage 2** fit PFU using plaque assay data, and **Stage 3** incorporated assumptions and parameter estimates from **Stages 1** and **2** to fit qPCR and PFU data together. At each stage, we compared competing models and estimated the necessary parameters **(Fig 2 A-C, E-G)**. We took this approach to allow the sequential incorporation of mechanistic conclusions and to limit model parameter identifiability issues. For each stage, we calculated Akaike Information Criterion (AIC) scores to determine the model that best fit the data with a minimal number of free parameters. The model with the lowest AIC score was selected.

In **Stage 1**, we fit competing models to qPCR data to estimate cell-associated (*V*_*Cell,DNA*_) and supernatant (*V*_*Sup,DNA*_) viral DNA dynamics from high and low MOI experiments **(Fig 1A, B)**. We estimated viral DNA production rate (*π*_*DNA*_), eclipse phase transition rate (*κ*) and mechanism (single versus multi-stage eclipse phase), death rate of infected cells (*δ*_*I*_), viral DNA egress parameters and mechanisms (density-dependent versus linear egress) and contribution of cell burst to viral DNA in the supernatant (*f*_*DNA*_). The estimated parameter values and preferred mechanisms (discussed in detail below) were carried forward to **Stages 2** and **3**. In **Stage 1**, The cell-free (*β*_*sup*_) and cell-associated (*β*_*cell*_) infectivity parameters were used only for fitting to low MOI data, since most cells were already infected at t=0 in the high MOI. Cell-free infectivity was estimated to be negligible (∼10^−11^) and removed from the model at this stage, suggesting that most of the viral spread following low MOI infection occurred via cell-to-cell propagation.

In **Stage 2**, we fit competing models to plaque assay data to estimate cell-associated (*V*_*Cell,PFU*_) and supernatant (*V*_*Sup,PFU*_) PFU dynamics from high MOI experiments **(Supplementary Fig S2)**. Since plaque assay data were only available for the high MOI experiment, where almost all cells are infected immediately after inoculation, the simplified model did not include the susceptible cell compartment or the infectivity (*β*) parameter (details in the **Materials and Methods, equations 2**). We used the same eclipse phase mechanism and death rate estimated in **Stage 1**. We estimated the PFU production rate (*π*_*PFU*_) and the mechanism (density-dependent versus linear) and rate of PFU egress in the supernatant. The release of PFU following cellular burst (*f*_*PFU*_), was estimated to be negligible and removed from the model. Preferred mechanisms and parameter values were carried forward to **Stage 3**.

The final **Stage 3** model showed an extremely close fit to qPCR and plaque assay data for cell-free and cell-associated virus following high MOI infection **(Fig 3A)**, and to qPCR data following low MOI infection **(Fig 3B)**, as well as projections of infectious viral infectious particle kinetics following low MOI infection **(Fig 3B). Table 1** shows a summary of the models tested at each stage with their lowest AIC score out of 6 assessment runs with different initial guesses of the parameters. Details about the mathematical equations, initial conditions, and fitting methods used at each stage are discussed in the **Materials and Methods** section, **Equations 3. Supplementary Table S2** includes all the model variations tested and their AIC scores, with the model selected highlighted in bold.

**Table 1.** Summary of tested models in each stage. The fixed and estimated parameters of each model, as well as their AIC scores, are listed. The AIC score is the lowest of the six assessment runs of each model, with different initial guesses for parameter values.

| Model | Fixed parameters | Estimated parameters | Lowest AIC |
| --- | --- | --- | --- |
| Stage 1, fitting to qPCR data |  |  |  |
| Linear egress rate | $\gamma = 0.04$ | $\log_{10} \beta_{Sup}, \log_{10} \beta_{Cell},$<br>$\kappa, \pi, D, \delta$ | 31.45 |
| Density-dependent egress rate | $\gamma = 0.04$<br>$\beta_{Sup} = 0$ | $\log_{10} \beta_{Cell},$<br>$\kappa, \pi, D, K, \delta$ | 8.55 |
| Density-dependent egress rate with burst | $\gamma = 0.04$<br>$\beta_{Sup} = 0$ | $\log_{10} \beta_{Cell},$<br>$f, \kappa, \pi, D, K, \delta$ | 6.65 |
| Stage 2, fitting to plaque assay data |  |  |  |
| Density-dependent egress rate | $\gamma = 0.04$<br>$\delta = 0.3$ | $\kappa, \pi, D, K$<br>$V_{Cell}(0), V_{Sup}(0)$ | -59.15 |
| Linear egress rate | $\gamma = 0.04$<br>$\delta = 0.28$ | $\kappa, \pi, D$ | -63.29 |
| Linear egress rate with burst | $\gamma = 0.04$<br>$\delta = 0.28$ | $f, \kappa, \pi, D$ | -70.82 |
| Stage 3, fitting to qPCR and plaque assay |  |  |  |
| Density-dependent egress rate with burst term of viral DNA | $\gamma_I = 0.04$<br>$\gamma_{NI} = 0$<br>$\delta = \delta_{PCR}$<br>$\pi_{PFU}, \pi_{DNA},$<br>$D_{PFU}, D_{DNA}, K, f_{DNA}$ | $\log_{10} \beta_{Cell},$<br>$\kappa$ | -25.29 |
| Density-dependent egress rate with burst term of viral DNA and conversion of viral DNA to PFU | $\gamma_I = 0.04$<br>$\gamma_{NI} = 0$<br>$\delta = \delta_{PCR}$<br>$\pi_{DNA},$<br>$D_{PFU}, D_{DNA}, K, f_{DNA}$ | $\log_{10} \beta_{Cell}, \alpha$ | -26.00 |
| Density-dependent egress rate with burst term of viral DNA with two stages of production | $\gamma_I = 0.04$<br>$\gamma_{NI} = 0$<br>$\delta = \delta_{PCR}$<br>$\kappa, \pi_{PFU}, \pi_{DNA},$<br>$D_{PFU}, D_{DNA}, K, f_{DNA}$ | $\log_{10} \beta_{Cell}, \lambda$ | -30.54 |

**Figure 3.**
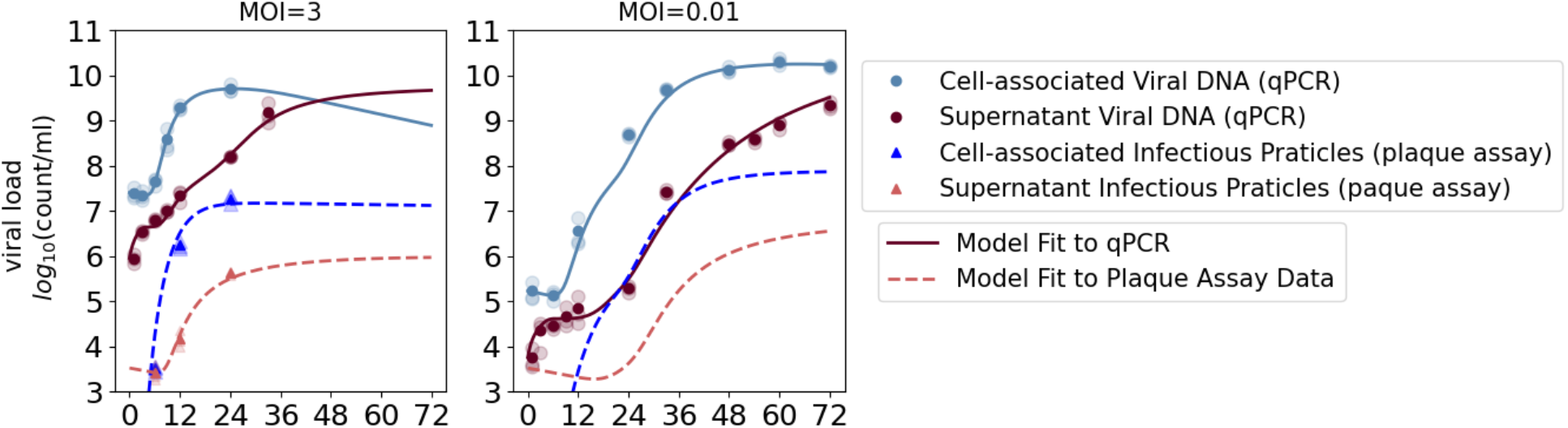
Optimized mathematical model recapitulating cell-associated and cell-free viral DNA and infectious particles in Vero cells following low and high MOI infection. The dark data points show the average viral load obtained from the three biological replicates at each time point. The light data points show the three replicates. The solid line and dashed line show the model fit to qPCR and plaque assay data, respectively. High MOI (left panel) and low MOI (right panel) experiments are shown. Plaque assay data was not obtained from low MOI experiments and only model projections are shown in this case.

### Normally-distributed eclipse phase lifespan in infected Vero cells

The eclipse phase describes the pre-productive phase of infection, following viral entry and before cells start the production of new viruses. The time spent in the eclipse phase was described by the Erlang (*κ, n*) distribution (a special case of the Gamma distribution with an integer shape parameter), where the rate parameter *κ* represents the rate of transition from one eclipse stage to the next, and the shape parameter *n* is the number of compartments. A low number of compartments translated to an exponentially distributed lifespan with a high variability between cells in the duration of eclipse, whereas a high number of compartments resulted in a normally distributed lifespan with a more uniform duration of the eclipse phase. In **Stage 1**, we compared competing models with different numbers of eclipse compartments (**Fig 2A and E**). The best model was selected based on the lowest AIC score and included 15 eclipse compartments **(Fig 4A)**. The transition rate estimated for the high and low MOI data with n=15 was *κ* =1.37 and 0.82, translating to an average time of 10.9 and 18.3 hours spent in the pre-productive eclipse phase, respectively. For n=15, the distribution was almost symmetrical around the mean, indicating that ∼90% of cells spent 7-16 hours (at high MOI) and 11-27 hours (at low MOI) in the eclipse phase (**Fig 4B**). However, the first cells started viral DNA production as early as 2.5 hours after infection at high MOI, and after 6.5 hours at low MOI.

**Figure 4.**
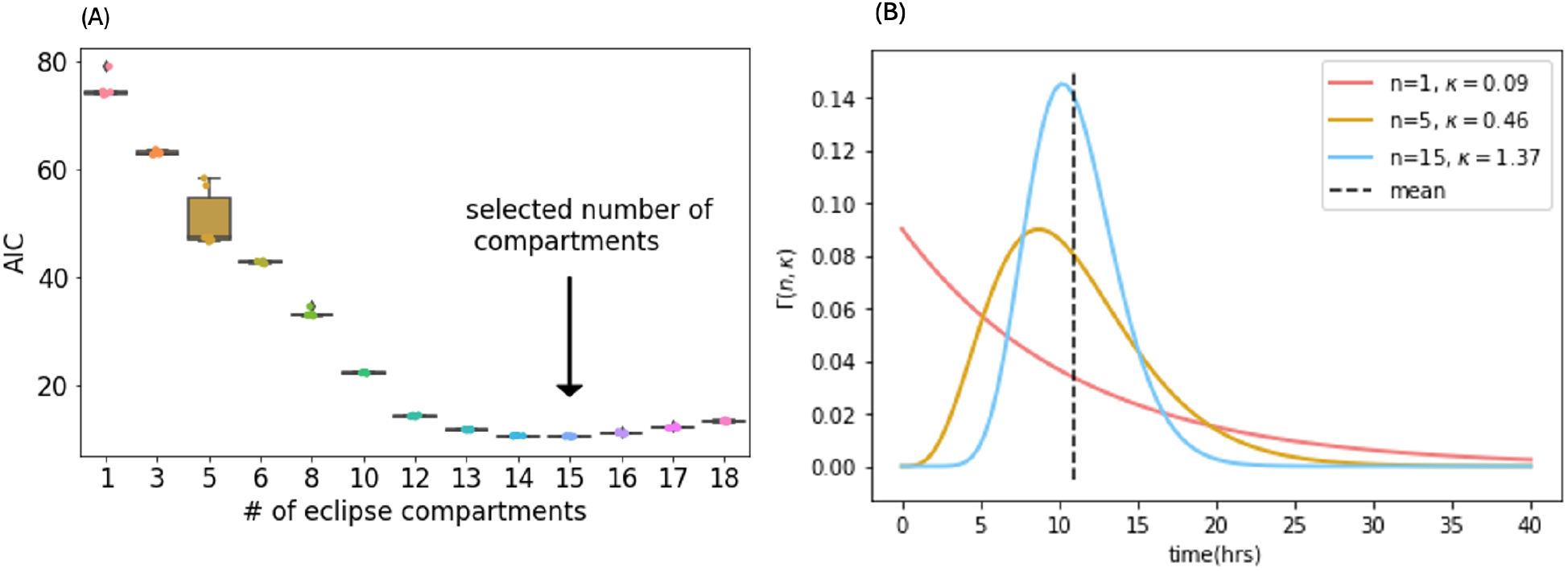
Infected cells enter an eclipse phase with a symmetrically distributed lifespan. (A) AIC scores of fit to qPCR data versus the number of stages of eclipse phase (n), with six different model runs for each n, each run starting with different initial parameter values. The model with 15 stages of the eclipse phase had the lowest AIC score. (B) Gamma distribution with different rates and shape parameters *κ, n*, with the same mean transition time. Distributions with large shape parameter *n* appeared symmetrically distributed around the mean. For n=15, the *κ* = 1.37 was estimated for high MOI experiments.

In **Stage 2**, we used the same 15 eclipse phase compartments estimated in **Stage 1**. The transition rate from eclipse to productive phase was estimated to be lower for PFU than viral DNA, indicating a delay between the production of viral DNA and infectious particles. To account for this delay, the parameter *λ* was introduced in **Stage 3**, representing the transition rate from viral DNA to PFU production. This model obtained the lowest AIC score. *λ* was estimated as *λ* = 0.53 for both high and low MOI. This indicated that, on average, infected cells started producing infectious particles 1.9 hours after viral DNA production had initiated.

### DNA and PFU exit from Vero cells

Viral egress was modeled as a diffusion rate of viral DNA and PFU from the cells to the supernatant. For viral DNA in **Stage 1**, a density-dependent diffusion rate in the form of 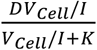 performed better than a constant diffusion rate (**Supplementary Fig S1**). This observation implied that the rate of viral DNA diffusion saturated with higher levels of viral DNA in infected cells over time.

In **Stage 2**, we compared competing models with density-dependent versus constant diffusion rates for PFU egress. The threshold parameter, *K*, in the density-dependent egress rate term, was estimated to be extremely small, suggesting that infectious particles egressed out of the cells at a constant rate without density dependence or saturation. While the fits for the two choices of egress rate were identical, the model with a constant rate had a lower AIC and was selected. **Supplementary Fig S2** shows the model fit to plaque assay data.

In addition, in **Stage 1**, we included a burst of infected cells releasing HSV DNA into the supernatant (*f*_*DNA*_) after cell death, which further improved the fit (**Supplementary Table S2)**. Interestingly, the burst size of the infectious particles (*f*_*PFU*_) was estimated to be minuscule in **Stage 2** and removed from the model. This suggested that nearly all infectious particles detected in the supernatant were due to linear egress out of the cell.

In **Stage 3**, based on the results of the first two stages, density-dependent egress rate and constant egress rates were assumed for *V*_*Cell,DNA*_ and *V*_*Cell,PFU*_, respectively.

### HSV infectivity estimates

Our final optimized **Stage 3** model **(Fig 3)** allowed us to quantify each step of Vero cell infection following both high and low MOI infection. Following viral entry, cells spent an average of 11 and 18 hours in the pre-productive eclipse phase, in high and low MOI conditions, respectively **(Fig 5B)**. Following high MOI infection, 95% of susceptible cells immediately entered the eclipse phase **(Fig 5A, B)** and infection of the remaining susceptible cells initiated ∼6.5 hours later. Secondary infection occurred extremely rapidly with complete depletion of susceptible cells over 1.5 hours, suggesting efficient cell-to-cell spread **(Fig 5A)**. Ater low MOI infection, 1% of cells were initially infected and infection of the remaining susceptible cells initiated ∼13 hours later. Remarkably, following low MOI infection, nearly all cells (∼99%) were in eclipse at around 18 hours **(Fig 5B)**. This suggested that the small proportion of cells (∼0.15%) that entered the productive stage early (before 18 hours) were responsible for the secondary infection of all remaining susceptible cells.

**Figure 5.**
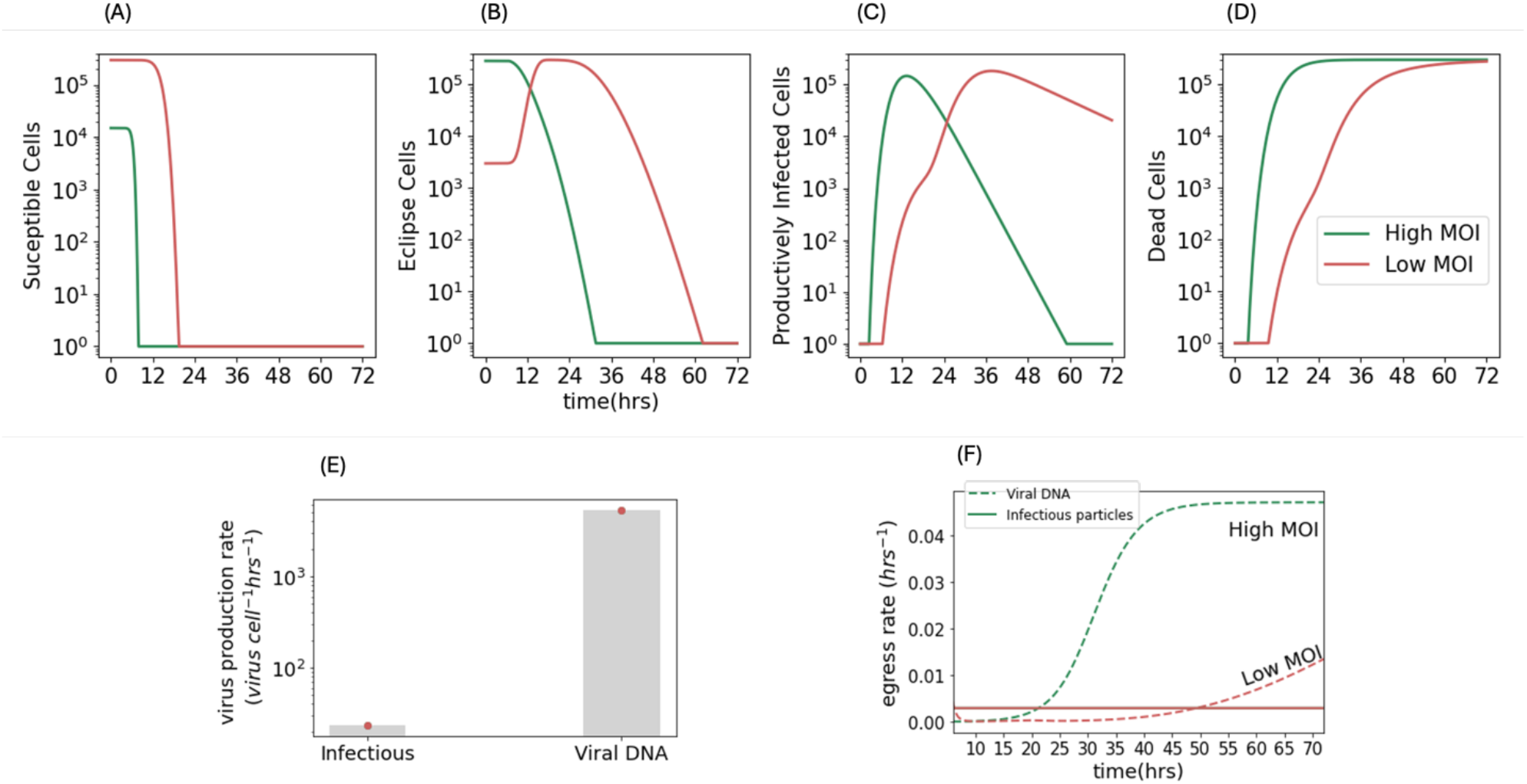
Faster production and egress of viral DNA relative infectious virus. (A-D) Dynamics of different model compartments in high and low MOI experiments. (E) Production rate (*π*) of viral DNA and infectious particles. (F) Egress rate of viral DNA and infectious particles. Once viral production begins after around 6 hours, infectious virions diffused out of the cells at a constant rate (solid lines, high and low MOI lines are overlapping), while viral DNA diffused at a density-dependent rate as a function of its population inside the cell

Under both low and high MOI conditions, infected cells started producing infectious particles an average of 1.9 hours after producing HSV DNA. The earliest cells entered the productive phase 2.5 and 6.5 hours following high and low MOI infection, respectively (**Fig 5C**). Less than 10 productive cells accounted for the observed expansion of HSV DNA within 12 hours of low MOI infection **(Fig 1B, 3B)**.

Infected cells also died at a faster rate following high MOI infection **(Fig 5C,D)**, with a rate of 0.29 and 0.072 deaths per hour at high and low MOI, respectively. This corresponded to a half-life of ∼3.5 and ∼14 hours after the start of the productive phase, or a total half-life of 14.5 and ∼32 hours after viral entry, for high and low MOI, respectively.

### Faster production and egress of viral DNA versus infectious particles in Vero cells

For every infectious particle produced, an average of 224 viral DNA molecules were produced (**Fig 5E**). While infectious virions egressed at a constant rate, the rate at which HSV DNA exited cells depended on HSV DNA concentration in cells. As the concentration of HSV DNA increased, the rate of egress also grew before ultimately saturating. Viral DNA eventually exited infected cells about 15 times faster than infectious virions (**Fig 5F)**. Viral DNA egress rate was higher following high versus low MOI infection (**Fig 5F**). In our infection system lacking immune responses, infectious particles decayed slowly while HSV DNA did not.

### Optimal mathematical model selection based on fit to HSV-1 kinetic data following N2A cell infection

Our experimental data showed that HSV-1 infection progressed differently between Vero and N2A cells (**Fig 1**). We therefore tested whether our mathematical models could account for these differences. Since plaque assay data was not available for N2A cells, a simpler one-stage model was used to recapitulate viral dynamics measured by qPCR in N2A cells **(Fig 6)**. After testing multiple competing models, the best model again included susceptible cells infected through cell-to-cell contact with negligible infectivity of cell-free virions. Once infected, cells entered a pre-productive eclipse phase before producing viruses. The time spent in the eclipse phase followed the Erlang distribution with only one compartment **(Fig 7)**. Productively infected cells produced cell-associated viruses at the rate of *π*. Low cell death was observed experimentally during N2A infection. In agreement, the model estimated the cell death rate of infected cells to be very low and it was set to zero. Cell-associated HSV DNA egressed into the supernatant at a constant rate *D*, and cell-free viral DNA degraded at the rate *γ* (**Fig 6A**).

**Figure 6.**
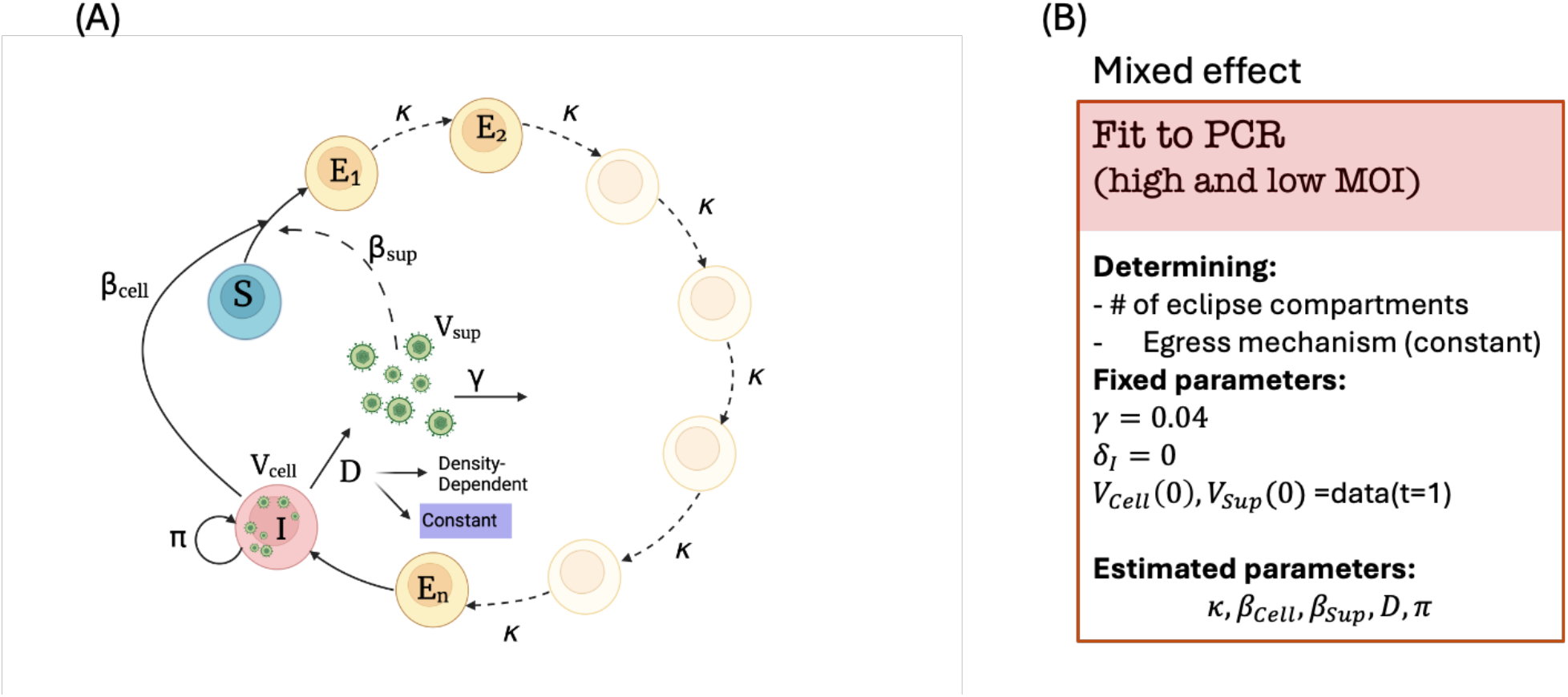
Mathematical model fit to N2A data and the modeling approach. (A) Schematics of the mathematical model with susceptible cells (*S*), Cells in the eclipse phase (*E*_*i*=1,*…,n*_), and productively infected cells (*I*), which produce viral particles (infectious and non-infectious) at rates *π*. Viral particles diffuse to supernatant at a constant or density-dependent diffusion rate. The susceptible cells are infected through cell-to-cell contact or by free virions in the supernatant at rates *β*_*Cell*_ and *β*_*Sup*_. Dashed arrows show the mechanism considered in the model and then removed due to negligible estimated parameter value. (B) Fixed and estimated parameters when fitting the model qPCR data.

**Figure 7.**
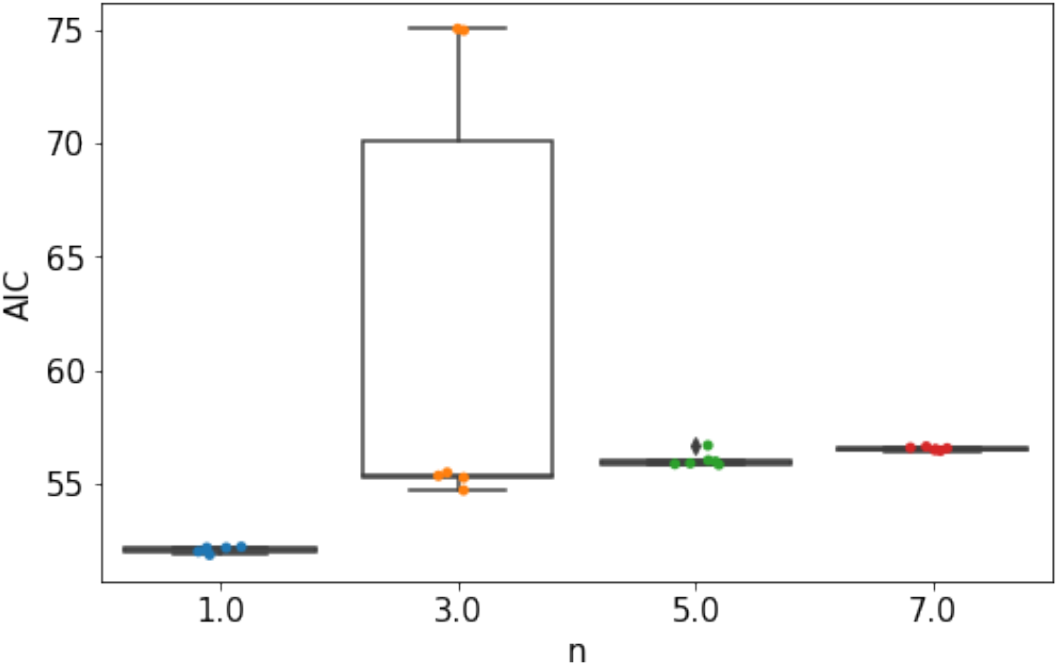
The model with one eclipse phase recapitulates N2A data with the lowest AIC score. AIC scores of fit to N2 data versus the number of stages of eclipse phase (n)

The best model fit to cell-associated and cell-free viral DNA levels in N2A cells had only one stage of the eclipse phase (n=1, **Fig 7**), implying an exponentially-distributed duration of eclipse, and a linear egress rate. A density-dependent egress rate was tested, but the threshold parameter, K, was estimated to be negligible and not identifiable. Model data fit was good, even though some non-monotonicity in the data following low MOI infection was not captured **(Fig 8)**.

**Figure 8.**
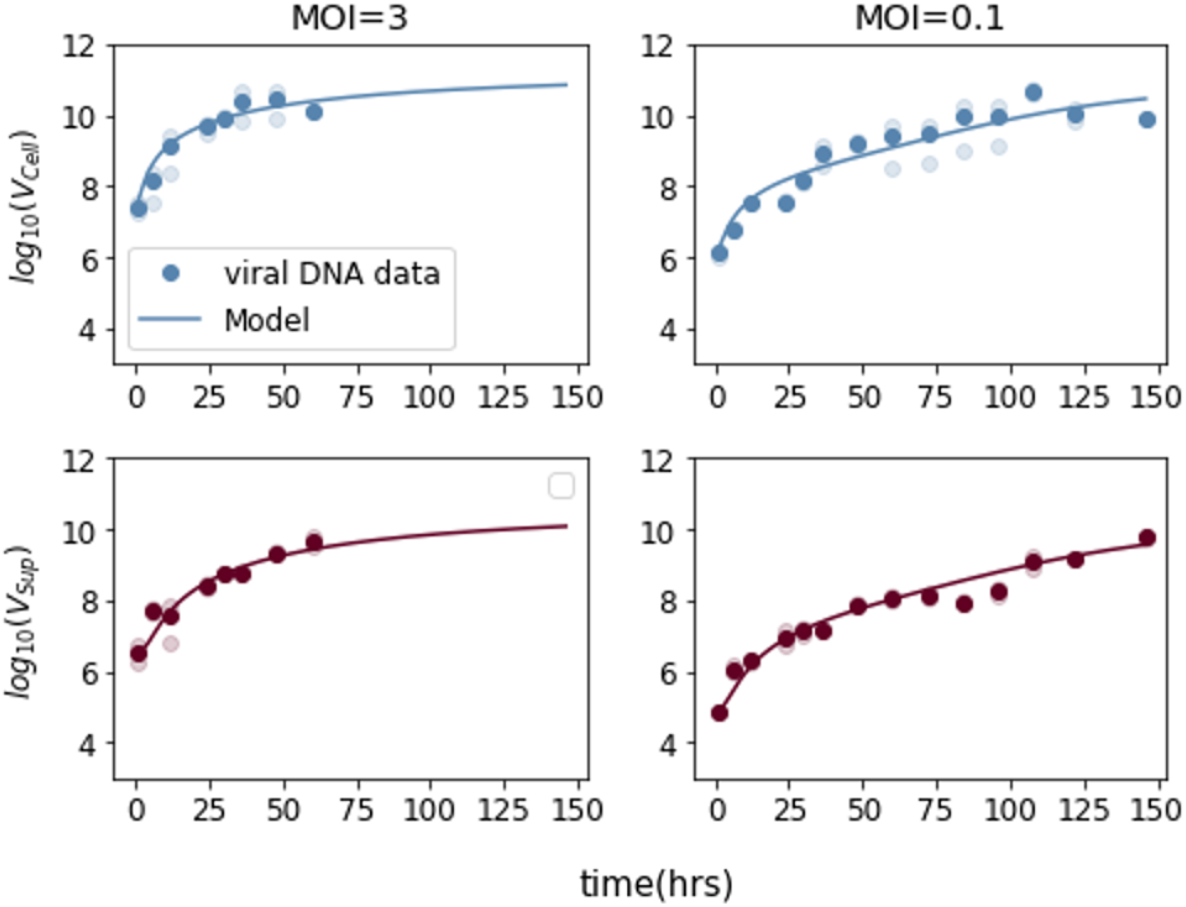
Model fits to N2A PCR data. Constant diffusion rate and one stage of the eclipse phase best described the data. The death rate of the infected cells was estimated to be negligible and set to zero *γ* = 0.04, and the first data point at t=1 was taken as the initial values of *V*_*Cell*_ and *V*_*Sup*_. The model was fitted to the average data (dark dots). The light dots show the individual replicates at each time point

### Differences in eclipse phase, viral production rate, and diffusion dynamics between Vero and N2A cells

According to our models, the distribution and average time spent in the pre-productive phase, virus production rate, and egress mechanisms differed significantly between Vero and N2A cells (**Fig 9**). Vero cells spent an average of 11 or 18 hours in the pre-productive phase (depending on MOI), with a low variability (Standard deviation SD=3.65 hours) among infected cells. By contrast, N2A cells had a much longer average eclipse phase (∼ 41.6 hours) and higher variability between cells (SD=41.6 hours). In the high MOI experiment, most neurons were infected after inoculation (**Fig 10A**). A small number of neurons became productive almost immediately after viral entry, whereas many neurons remained non-productive a week after infection **(Fig 10B, C)**. The time spent in the eclipse phase for Vero cells appeared symmetrically distributed around the mean, while for the N2A cells, the distribution was exponential, causing a large variability in the transition rate (**Fig 9A**).

**Figure 9.**
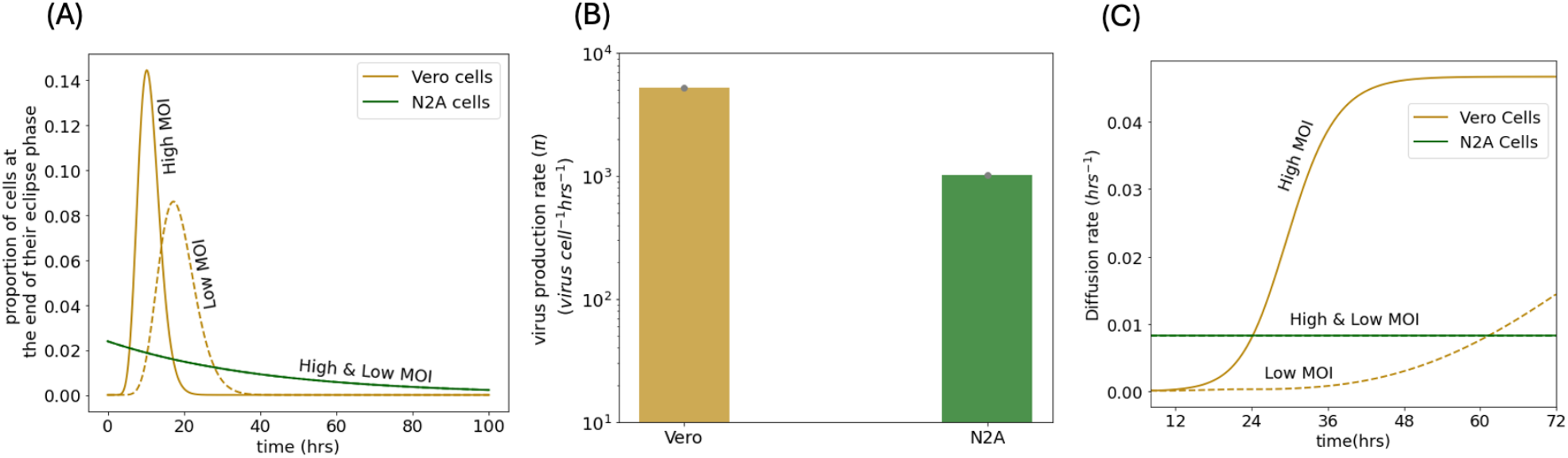
Different HSV DNA dynamics in Vero and N2A cells. (A) Distribution of the duration of the eclipse phase for Vero and N2A cells estimated by the model fit to qPCR data. In Vero cells, the distribution was close to normal and cells spent an average of 14 hours in the pre-productive phase after being infected. In N2A cells, the exponential distribution had a long tail and cells spent a highly variable and on average much longer time in the pre-productive phase. (B)Rate of virus production in Vero cells and N2A cells. (C) Diffusion rate over time, with either a density-dependent rate (Vero cells), or constant rate (N2A cells)

**Figure 10.**
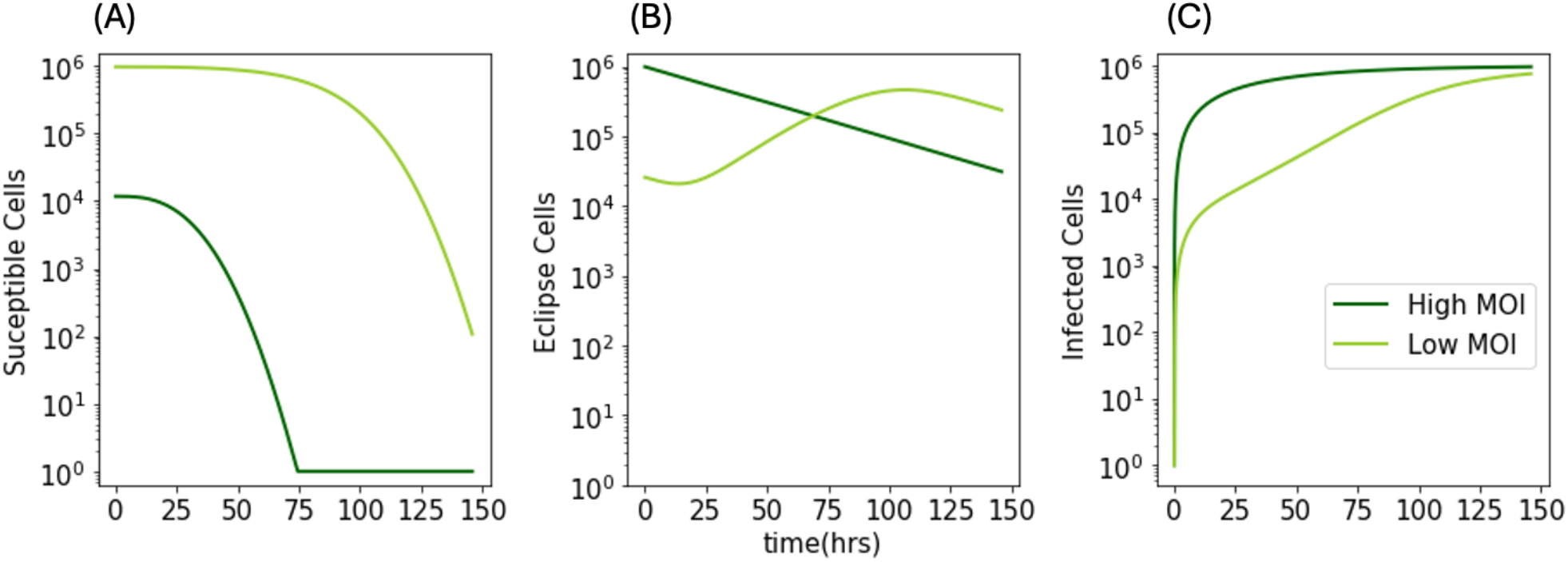
Dynamics of different model compartments of N2A cells in high and low MOI.

The estimated HSV DNA production rate in Vero cells was around 5-fold higher than in the N2A cells (**Figure 9B**). Our model predicted that in N2A cells, HSV DNA egressed into the supernatant at a constant rate (**Figure 9C**). Due to the immediate appearance of the productively infected cells, a slow but steady egress of viruses started immediately after infection. By contrast, in Vero cells, egress started after ∼2.5 and ∼6.5 hours after infection for the high and low MOI experiments, respectively, and accelerated thereafter.

## Discussion

We applied mathematical models to highly detailed viral kinetic data, which revealed differences in cell-associated and cell-free levels of viral DNA and PFU following high and low MOI infection in epithelial and neuronal cells. These models arrived at the following conclusions: 1) the average eclipse phase was longer in Vero cells than expected, with a small percentage of early-producing cells contributing to production of HSV DNA and PFU early during infection; 2) the rate of viral DNA production was much more rapid than the production of infectious particles in Vero cells; 3) HSV DNA exited cells faster than infectious particles from Vero cells and did so in a density-dependent saturating fashion; 4) in Vero cells, high MOI conditions led to a shorter eclipse phase, shorter time to viral egress and more rapid lysis of infected cells relative to low MOI conditions; 5) cell-to-cell spread of HSV PFU likely accounted for the majority of new cellular infections in both cell types, as the infectivity of cell-free viruses was negligible; 6) the eclipse phase of infection had much higher variability in neuronal cells than epithelial cells and was on average considerably longer; 7) HSV DNA production was more rapid in epithelial cells than neurons; 8) HSV DNA egress rate was dependent on the density of the cell-associated viruses in infected cells in epithelial cells but not in neuronal cells. Overall, these conclusions offer a very detailed analysis of the viral replication and spread cycle in both cell types under low and high MOI conditions in culture.

Each of these conclusions warrants further mechanistic consideration. The normal distribution of eclipse phase duration in epithelial cells suggests that the fate of most of these cells is only somewhat variable following infection. Only a small proportion of cells started producing viruses within a few hours of infection, and a small population also remained in the eclipse phase for more than 20 hours. The rapid rate of egress and subsequent spread suggested that early productive infection in a few cells may have an outsize role in facilitating infection of surrounding cells under low MOI conditions.

By contrast, the exponential distribution and prolonged of eclipse phase in neurons was consistent with a predisposition towards latency during which DNA amplification is limited. In vivo, HSV establishes long-term latency in neurons, indicating that neurons can efficiently restrict productive infection. Latency may be established immediately after entry or may be accompanied by a short burst of HSV DNA replication or even productive infection^24,25^. Our model accurately recapitulated that characteristic. In our cell culture experiment, due to consistently high concentration of viruses in the dish, all cells eventually became productively infected. Yet, our model indicated that some neurons remained in the pre-productive phase for days. We hypothesize that this extended eclipse phase reflects a predisposition toward long-term latency. In vivo, immune or other mechanisms could further tip the balance and push HSV toward a latent state.

The more rapid production of viral DNA relative to PFU in Vero cells was expected. Previous studies have reported DNA to PFU ratios in the range of 10 to 100 in supernatant^26–28^. Our experimental data was on the higher side of that range, with a DNA to PFU ratio of 368 and 273 for cell-free and cell-associated particles, respectively, at 24 hours at high MOI. Consistent with our experimental data, our model predicted that an average of 224 viral DNA molecules were produced for every infectious particle, and the production of the infectious particles started with a 1.9-hour delay relative to the first DNA production. It would be a useful next step to provide a quantitative framework on the steps that occur between DNA amplification by the viral polymerase and packaging of genomically intact, fully infectious viral particles. Accentuating these bottlenecks may be a target for future possible therapies such as capsid inhibitors^29–32^. It is relatively rare to obtain plaque assay data in humans as viral culture is laborious and insensitive. Our results may therefore allow coarse projection of PFU levels in human infection when this data is lacking. We recently demonstrated for SARS-CoV-2 that two drugs with equivalent potency but different mechanisms, may impact the clearance kinetics of viral genomic levels quite differently despite having a comparable impact on infectious virus^18,33^. Our model suggests this is a plausible consideration for HSV as well.

The dynamics of viral egress are rarely considered in mathematical models, nor is the infectivity of cell-associated versus cell-free viruses. Our results demonstrated that cell-associated viruses account for nearly all infectivity following low MOI infection in epithelial cells. The mechanisms underlying this are likely related to spatial features of infection. Cell-to-cell spread of cell-associated viruses occurs through tight junctions^34,35^. However, our prior agent-based models demonstrate that this need not be restricted to adjacent cells. For HSV, cell-associated viruses can likely passage through tight junctions to cells that are multiple contacts away, allowing for a single infected cell to infect dozens of surrounding cells. This is likely the only plausible mechanism to explain how human HSV ulcers accrue hundreds of thousands of infected epithelial cells in less than a day^16^. Supporting this hypothesis, we found that a very small proportion of early productive cells could infect all remaining susceptible cells very quickly, by cell-to-cell spread (**Fig 5**). This suggested that a single infected cell could infect cells further away than its immediate neighbors.

Viral egress via the apical cell surface is important to consider for measuring cell-free viruses, which are measured in human infections in plasma and from mucosal swabs^36–38^. Indeed, while cell-free viruses do not appear to play an important role in viral spread within a plaque, they are likely vital for sexual and oral transmission of HSV between people^39^. The results of our modeling of HSV-2 genital infection are broadly consistent with this theme. In those models, we also projected much higher cell-associated HSV viral loads relative to those outside of the cell^13,14,17,21^. Yet, we predicted that cell-free HSV is responsible for the seeding of new micro-environments of infection, which is one reason that HSV lesions are defined by the formation of sequential crops of ulcers, and highly variable spatial viral loads^40^.

A final consideration is the more rapid dynamics of eclipse phase, viral egress and cell death following high versus low MOI infection. We were surprised by this finding because even under a low MOI=0.01 scenario, the 99% of infected cells that are not initially infected are subsequently exposed to a very high number of HSV particle forming units, which is similar to what all cells experience immediately after high MOI inoculation. A key difference pertains to the fact that high MOI conditions provide immediate, synchronized exposure whereas viral spread leads to more protracted exposure. Under high MOI infection, multiple cells simultaneously experience cytopathic effect which more quickly distorts the architecture of surrounding cells and appears to dramatically impact their survival. Our opinion is that low MOI infection is a much closer proxy of human HSV reactivation in which Poisson statistics suggest that infection within a given micro-environment is likely to start with one virus initiating infection in a single cell^4,5^. At a more basic level, high MOI infection neglects radial cell-to-cell spread which is central to in vitro plaque and in vivo HSV ulcer expansion.

The major limitation of our work was that the in vitro system did not capture other critical features of infection in humans, particularly immune responses. This was readily evident in our model as HSV DNA decay was not needed to recapitulate the virologic data, while HSV DNA contracts extremely rapidly during human infection^17,41^. Viral levels plateaued rather than decreased after peaking, as is sometimes the case in human infection in severely immunocompromised individuals^42^. Another limitation of our differential equation system is that it does not capture spatial features of infection. Using agent-based models applied to viral data in spreading plaques, we previously demonstrated that target cell limitation occurred extremely early during infection and that innate immunity likely relates to spreading cytokines from initially infected cells^16^, both of which are not captured by our model or our data. Despite this limitation, we feel there is a high utility in isolating the specifics of viral replication, egress, and infection without considering the multifactorial effects of immune responses. Finally, we do not interrogate the numerous steps of HSV replication and gene expression in detail, despite the fact that these clearly relate to the output of HSV DNA and infectious virus. This would be an interesting pursuit but would require single-cell expression data and modeling.

In conclusion, we developed an unprecedently detailed model of HSV replication and spread which accounts for discordance between HSV DNA and infectious virus production, heterogeneity in viral eclipse phase, kinetics of egress, differential infectivity of cell-associated and cell-free virus, and different behaviors between cells which are predisposed to lytic versus latent infection.

## Materials and Methods

### Cells and viruses

African green monkey epithelial Vero cells and murine neuroblastoma N2A cells were obtained from the ATCC and cultured in DMEM (Corning, Corning, NY, USA) supplemented with 10% FBS (Sigma-Aldrich, St-Louis, MO, USA). Cells were maintained at 37 °C in a 5% CO_2_ humidified incubator and frequently tested negative for mycoplasma contamination.

For HSV-1 infection, we used an HSV-1 virus from strain 17+ expressing YFP from the US1/2 locus, which was described previously^43^. Infections were conducted in 12 well plates. At the time of infection, confluent wells contained approximately 3 × 10^5^ Vero cells, or 1 million N2A cells. Cells were infected at the indicated MOI for 1 hour before removing and replacing the inoculum with 1mL of fresh media. At each time point, 50*μ*L of supernatant was collected in a vial containing 1mL of digestion buffer (KCL, Tris HCl pH8.0, EDTA, Igepal CA-630) and stored at 4°C before DNA extraction. To collect cells, the supernatant was removed and cells were washed once with PBS. Cells were either mechanically or enzymatically detached from the plate and resuspended in 1mL of PBS. 50uL of the suspension was then collected in 1mL of digestion buffer or directly analyzed by plaque assay.

Plaque assays were conducted in Vero cells. Confluent Vero cells in 24-well plates were incubated for 1 h with 100uL of inoculum and 10-fold serial dilutions. Cells were overlaid with 1mL of complete media containing 1% methylcellulose, prepared using DMEM powder (ThermoFisher, USA) and Methylcellulose (Sigma-Aldrich, USA). After two or three days, fluorescent plaques expressing YFP were acquired with an EVOS automated microscope (ThermoFisher, USA) and counted using with ImageJ (v2.14.0/1.54f).

For qPCR, DNA was extracted from 200 μl of digestion buffer using QiaAmp 96 DNA Blood Kits (Qiagen, Germantown, MD, USA) and eluted into 100 μl AE buffer (Qiagen, Germantown, MD, USA). 10 μL of eluted DNA was used to set up 30 μL real-time Taqman quantitative PCR reactions, using QuantiTect multiplex PCR mix (Qiagen, Germantown, MD, USA), using the following PCR cycling conditions: 1 cycle at 50°C for 2 minutes, 1 cycle at 95°C for 15 minutes, and 45 cycles of 94°C for 1 minute and 60°C for 1 minute. Exo internal control was spiked into each PCR reaction to monitor inhibition. A negative result was accepted only if the internal control was positive with a cycle threshold (CT) within 3 cycles of the Exo CT of no template controls. Primers and probes have been described previously^44^ and recognized HSV *gB* (Fwd: CCGTCAGCACCTTCATCGA, Rev: CGCTGGACCTCCGTGTAGTC, Probe: FAM-CCACGAGATCAAGGACAGCGGCC).

qPCR and plaque assay data used for modeling and included in **Supplementary Table S5**.

## Mathematical models

The model to recapitulate viral DNA and plaque assay data in the Vero cell experiments was developed in three stages. In **Stage 1**, we developed a target-cell-limited model and parametrized it only using viral DNA (qPCR) data. We compared models with constant and density-dependent egress rates and with and without release of viruses to supernatant after the death of an infected cell (burst mechanism). For each of these models, we tested the model with 1, 3, and 5-18 stages of eclipse phase. Finally, we compared the model with cell-to-cell spread alone versus cell-to-cell spread and cell-free virus infection mechanisms. This resulted in 50 competing models. In **Stage 2**, the model was parametrized using high MOI only plaque assay data. We compared three competing models with constant and density-dependent egress rates, and with and without the burst mechanism. In **Stage 3**, we combined the model features of the two previous stages and parameterized the model using both viral DNA and plaque assay data. We compared two models with and without a delay between the DNA and infectious particles production, and another assuming a fraction of DNA converts to infectious particles. All competing models are listed in **Supplementary Table S2**. Most of the estimated parameters from **Stages 1** and **2** were fixed in stage three to help with parameter identifiability. **Supplementary Table S3** includes all the parameters estimated in each stage. Details about the model structure, initial conditions, and fixed parameters for each stage of model development are discussed below.

For the N2A cell experiments, we tested multiple target cell limited models with cell-associated and supernatant virus compartments with 1, 3, 5, and 7 stages of the eclipse phase. We also compared different egress rates and cell-to-cell and cell-free mechanisms of viral spread. All the tested models are listed in the **Supplementary Table S2. Supplementary Table S4** includes the parameters estimated for the N2a model.

The best model was selected based on the lowest AIC score. The AIC score, defined as −2 ln(*L*) + 2*m*, is a statistical score used to determine which model fits the data better with a minimal number of parameters^45^. AIC is rewarded by the goodness of fit (the first term, the log-likelihood of the model generating the observed data) and penalized by the complexity of the model (the second term, with m equals the number of parameters in the model).

### Viral dynamics model recapitulating viral DNA data in Vero cells (stage 1)

The winning model in **Stage 1** was a target cell limited model that includes susceptible cells (S), n=15 stages of eclipse phase (E_i=1,…,n_), productively infected cells (I), cell-associated viruses (V_Cell,DNA_), and the viral load in the supernatant (V_Sup,DNA_). The susceptible cells are infected through cell-to-cell contact at an average rate of *β*_*Cell*_*V*_*Cell,DNA*_. The rate of infection by free virions present in the supernatant (*β*_*Sup*_) was estimated to be very small and therefore removed from the model. The infected cells enter an eclipse phase with a mean transit time of *n/κ* after which they start producing viruses at rate *π*_*DNA*_. Cell-associated viruses egress out of the cells into supernatant at the rate 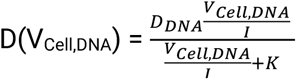, where D_DNA_ is the maximum egress rate and K is the threshold viral load per infected cell for which the egress rate is D_DNA_/2. The free viral particles clear at the rate *γ*. Viral particles in the cell are released to the supernatant after the burst of infected cells with burst size f_DNA_. Writing the model as a set of ordinary differential equations, we have,

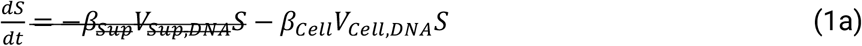

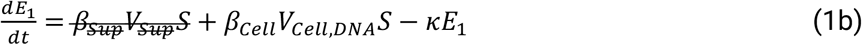

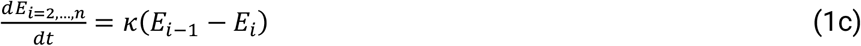

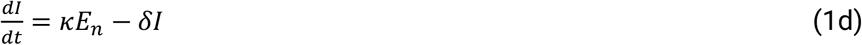

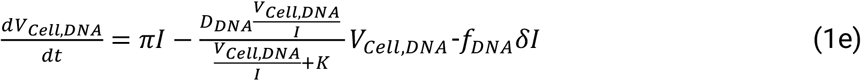

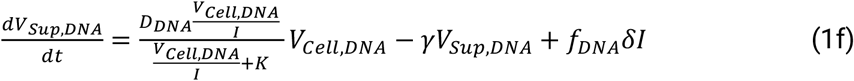

For the initial conditions of susceptible and infected cells (S and E1), we use the Poisson distribution to calculate the probability of one or more viruses entering a susceptible cell given MOI. The number of susceptible cells and infected cells in the eclipse phase at the beginning of the experiment is given by *S*_0_ = *N* exp(−*MOI*) and *E*_1,0_ = *N*(1 − exp(−*MOI*)) respectively, where *N* = 3 × 10^5^ is the total number of cells in each assay, and exp(−*MOI*) is the probability of a cell remaining uninfected (no viruses entering the cell). We set *E*_*i*=2,*…,n*,0_ = *I*_0_ = 0 and first data point was used for *V*_*Cell*,DNA_(0) and *V*_*Sup*,DNA_(0).

To limit parameter identifiability issues, we fixed *γ* = 0.04.

### Viral dynamics model recapitulating plaque forming unit data in Vero cells (stage 2)

The plaque forming unit data alone doesn’t provide any information about the infectivity of the virus (*β*_*cell*_ and *β*_*sup*_) since the data was only collected from the high MOI experiments in which almost all the cells were infected after initial inoculation. Therefore, when modeling plaque forming unit data, we assume all the cells are in the first stage of the eclipse phase at the beginning of the experiment. The model with a constant egress rate fits plaque assay data with a lower AIC score (**Supplementary Table S2**).

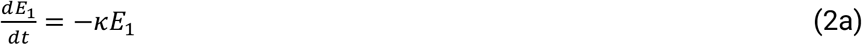

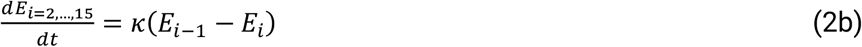

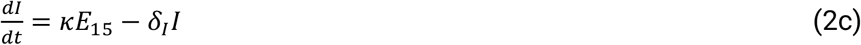

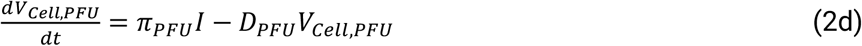

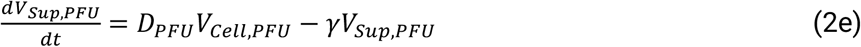

We used *V*_*Cell,PFU*_(0) = 0 and estimated *V*_*Sup,PFU*_(0) from the data. We fixed *γ* = 0.04, and *δ* = 0.28, as estimated by fitting the model to viral DNA data.

### Viral dynamics model recapitulating viral DNA and plaque assay data simultaneously (Stage 3)

In the combined model, cell-associated and supernatant virus compartments are each divided into infectious virions (*V*_*Cell,PFU*_ & *V*_*sup,PFU*_) and viral DNA (*V*_*Cell,DNA*_ & *V*_*sup,DNA*_). Infectious virions compartments of the model are fit to plaque assay data, while the sum of infectious virions and viral DNA are fit to viral DNA data. In this final version of the model, infectious virions egress at a constant rate (D_PFU_) while the rate at which viral DNA exits the cell with a density-dependent rate (*D*_*DNA*_(*V*_*Cell,DNA*_)). Once out of eclipse phase the infected cells begin producing viral DNA (I_DNA_). The production of infectious particles starts with an average delay defined by 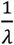. Infected cells producing both viral DNA and infectious particles are called I_PFU_ in the model.

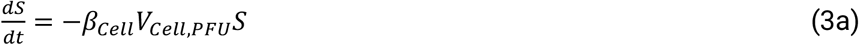

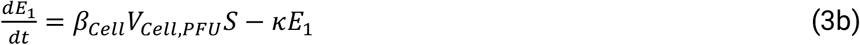

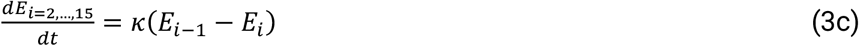

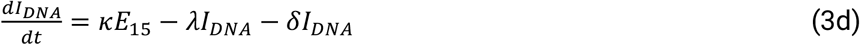

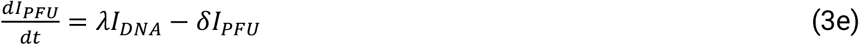

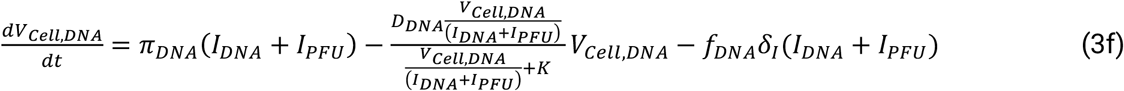

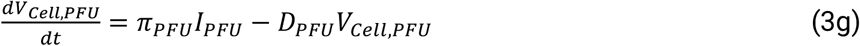

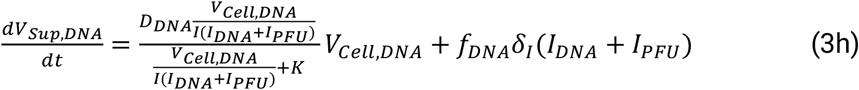

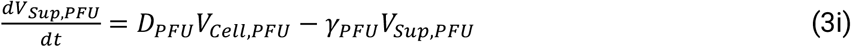

We used *V*_*Cell,PFU*_(0) = 0 and estimated *V*_*Sup,PFU*_(0). The initial values of the viral DNA particles (*V*_*Cell,DNA*_ & *V*_*sup,DNA*_) were calculated by deducting *V*_*Cell,PFU*_ and *V*_*sup,PFU*_ from the first viral DNA data point. To help with identifiability, we fixed the clearance rate of infectious virions *γ*_*I*_ = 0.04 and the clearance rate of viral DNA particles to zero, based on the observation that viral DNA remains stable and does not degrade in the time span of the experiment. All the parameters associated with the dynamics of viral DNA and infectious particles were imported from **Stage 1** and **Stage 2**, respectively. Only the cell-to-cell infectivity of infectious virions (*β*_*cell*_) and the delay between infectious virions and DNA production (*λ*) were estimated in this stage. The infectivity of free virions (*β*_*sup*_) and the burst size of infectious virions (*f*_*PFU*_) were set to zero based on the result of the first two stages of model fitting.

### Viral dynamics model recapitulating viral DNA data in N2A cells

The winning model that best recapitulates the dynamics of viral DNA data in the N2A cells is a target cell limited model that includes susceptible cells (S), one stage of eclipse phase (E_1_), productively infected cells (I), cell-associated viruses (V_Cell_), and the viral load in the supernatant (V_Sup_). The susceptible cells are infected through cell-to-cell contact at an average rate of *β*_*Cell*_*V*_*Cell*_. *β*_*Sup*_ was estimated to be very small and therefore removed from the model. The infected cells go through an eclipse phase with a mean transit time of 1*/κ* after which they start producing viruses at rate *π*. Cell-associated viruses egress out of the cells into supernatant at the constant rate D, and the free viral particles clear at the rate *γ*.

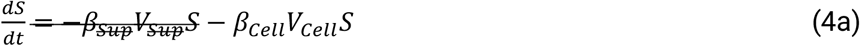

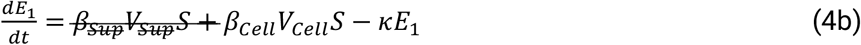

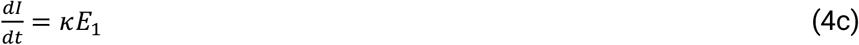

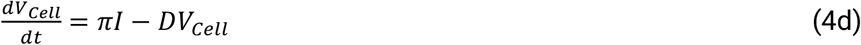

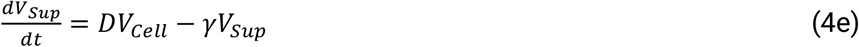

The initial value of susceptible cells (S) was estimated from the data using MOI as covariate, and the initial value of E1 was calculated as N-S(0), with N=10^6^, the total number of cells in each replicate. We set *I*_0_ = 0 and use the first data point as *V*_*Cell*,0_ and *V*_*Sup*,0_. Due to the long lifespan of N2A cells, we assumed infected cells did not die during the experiment. Therefore, no death term for the infected cells is considered in the model.

### Fitting and parameter estimation

Data used to parametrize the model was averaged over the three replicates at each time point. For the model fit to the Vero cell and N2A cell viral DNA data and the combined model, we used the population mixed effect approach implemented in Monolix to estimate the parameters.

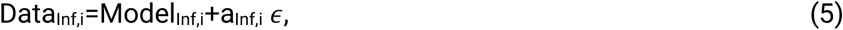

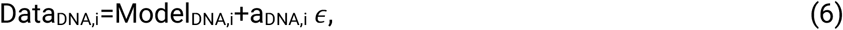

When fitting the models to viral DNA only and in the combined model, constant measurement error model was used in Monolix. In the combined model, error models are written as, where i=cell, sup, “a” is the error parameter, and *ϵ* is a normally distributed random number centered around zero with a standard deviation of one. Model_Inf,I_ represents the model compartments *V*_*Cell*.*PFU*_ and *V*_*sup*.*PFU*_, and Model_DNA,I_ refers to (*V*_*Cell*.*PFU*_ + *V*_*Cell,DNA*_) and (*V*_*sup*.*PFU*_ + *V*_*sup*.*DNA*_). When fitting the model to the plaque assay data, we used a simple least-square algorithm.

Values and model parameters for estimated viral dynamics quantities are indicated in **Supplementary Table S5**.

## Data availability

The data supporting the findings of this study are available within the paper and its Supplementary files.

## Code availability

All codes are available on GitHub at https://github.com/sEsmaeili/HSV-In-vitro-Modeling

## Acknowledgments

We thank members of the Schiffer and Jerome labs for technical and conceptual help. We thank Tracy K Santo at the University of Washington for running qPCR assays. This study was supported by NIH grant R21AI178255 and by a VIDD faculty initiative award from the Fred Hutch Cancer Center.

## Author contributions

Conceptualization: JTS, MW, SE. Methodology: JTS, SE, MW. Experimental design and data collection: MW. Computational modeling and software: SE. Analysis: SE, JTS, MW, DAS. Writing original draft: SE, JTS, MW. Writing review and editing: SE, JTS, MW, KRJ, DAS.

