## Supplementary Information for "Intracellular and extracellular dynamics of herpes simplex virus 1 DNA and infectious particles in epithelial and neuronal cells"

for

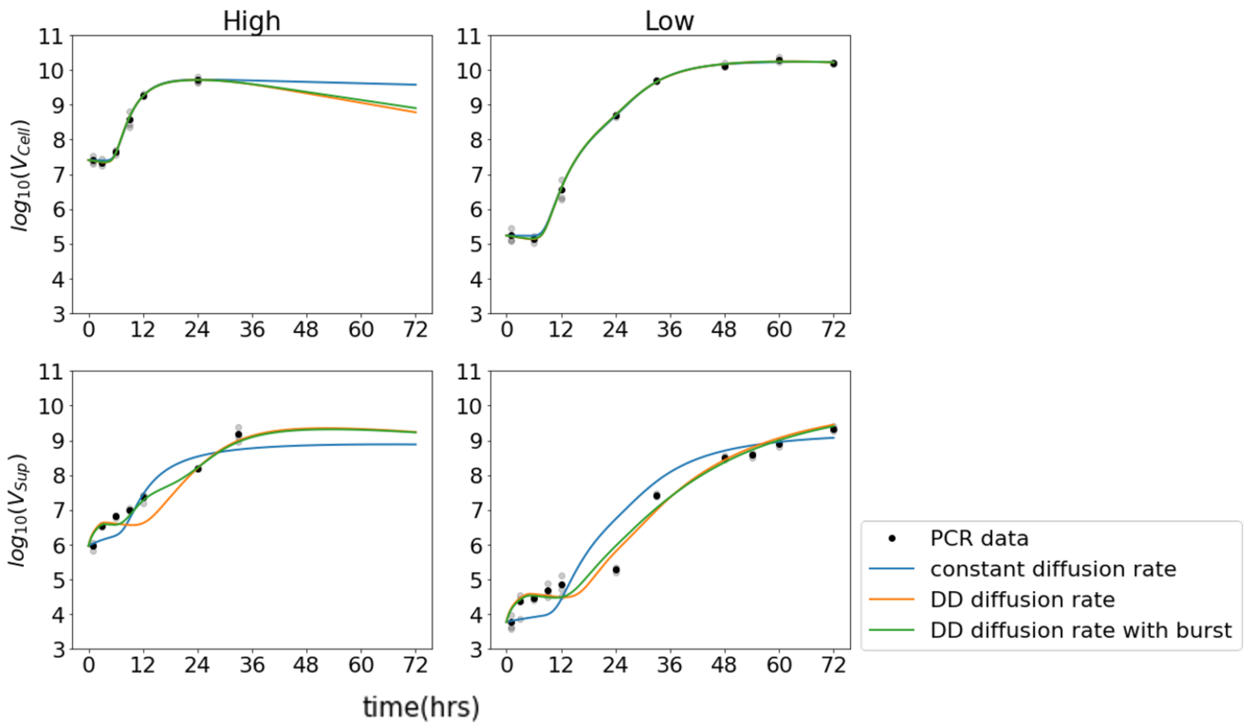


**Supplementary Figure S1. Density-dependent diffusion with burst recapitulates HSV DNA trajectories more accurately than linear diffusion.** Comparing model fits with density-dependent diffusion term with and without burst versus constant rate diffusion term to qPCR data ($\gamma$ is fixed to 0.04). The model with density-dependent diffusion term captures the delay in expansion of supernatant viruses in low MOI experiments better. Adding the burst term captures the initial increase in the supernatant viruses in the high MOI experiment (AIC = 6.65 vs 8.55 vs 12.44).

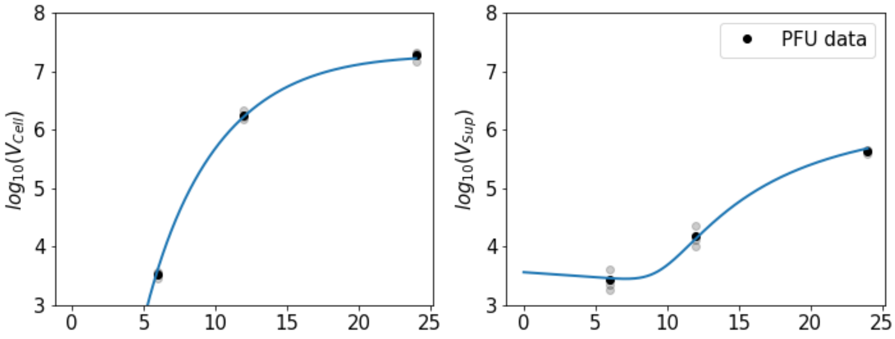


**Supplementary Figure S2. Constant egress rate recapitulates HSV pfu trajectories in Vero cells.** Model fit with linear egress term to PFU data ($\gamma = 0.04$, $\delta_I = 0.31$, $V_{Cell}(0) = 0$ ).

| Model | Fixed parameters | Estimated Parameters | Fit to | n | AIC |
|---|---|---|---|---|---|
| **Linear Diffusion**<br>$\frac{dS}{dt} = -\beta_{Sup}V_{Sup}S - \beta_{Cell}V_{Cell}S$<br>$\frac{dE_1}{dt} = \beta_{Sup}V_{Sup}S + \beta_{Cell}V_{Cell}S - \kappa E_1$<br>$\frac{dE_{i=2,\ldots,n}}{dt} = \kappa(E_{i-1} - E_i)$<br>$\frac{dI}{dt} = \kappa E_n - \delta I$<br>$\frac{dV_{Cell}}{dt} = \pi I - DV_{Cell}$<br>$\frac{dV_{Sup}}{dt} = DV_{Cell} - \gamma V_{Sup}$ | $\gamma = 0.04$ | $\log_{10}\beta_{Sup}$, $\log_{10}\beta_{Cell}$, $\kappa, \pi, D, \delta$ | Vero-PCR | 1 | 70.78<br>70.81<br>71.01<br>71.35<br>70.64<br>71.24 |
| | | | | 3 | 55.13<br>55.04<br>54.76<br>54.70<br>54.80<br>54.91 |
| | | | | 5 | 50.3<br>49.91<br>50.97<br>50.91<br>50.05<br>50.98 |
| | | | | 6 | 47.65<br>47.80<br>48.05<br>49.07<br>47.81<br>48.79 |
| | | | | 7 | 45.81<br>45.86<br>45.76<br>47.03<br>45.84<br>46.67 |
| | | | | 8 | 44.14<br>44.44<br>44.25<br>45.68<br>44.13<br>45.41 |
| | | | | 9 | 42.83<br>42.93<br>42.74<br>44.09<br>41.47<br>44.39 |
| | | | | 10 | 38.26<br>41.69<br>38.41<br>42.40<br>41.67<br>43.17 |
| | | | | 11 | 35.38<br>35.42 |

| | | | | | |
|---|---|---|---|---|---|
| | | | | | 35.63<br>35.79<br>35.58<br>41.70 |
| | | | | 12 | 33.26<br>33.46<br>33.53<br>40.79<br>33.50<br>34.77 |
| | | | | 13 | 32.17<br>32.29<br>32.38<br>40.71<br>32.17<br>40.60 |
| | | | | 14 | 31.6<br>31.55<br>31.68<br>31.63<br>31.56<br>39.92 |
| | | | | 15 | 31.45<br>31.48<br>31.53<br>31.47<br>31.53<br>31.51 |
| | | | | 16 | 31.64<br>31.76<br>31.83<br>31.63<br>31.74<br>39.02 |
| | | | | 17 | 32.05<br>32.16<br>32.16<br>31.96<br>32.06<br>32.24 |
| | | | | 18 | 32.82<br>32.69<br>32.79<br>32.57<br>32.60<br>38.44 |
| **Density-Dependent Diffusion (per cell)**<br>$\frac{dS}{dt} = -\beta_{Sup} V_{Sup} S - \beta_{Cell} V_{Cell} S$ | $\gamma = 0.04$ | $\log_{10} \beta_{Sup}$,<br>$\log_{10} \beta_{Cell}$,<br>$\kappa, \pi, D, K, \delta$ | Vero-PCR | 1 | 69.88<br>69.86<br>69.63<br>70.18<br>69.86 |

| | | | | | |
|---|---|---|---|---|---|
| $\frac{dE_1}{dt} = \beta_{Sup}V_{Sup}S + \beta_{Cell}V_{Cell}S - \kappa E_1$<br>$\frac{dE_{i=2,\dots,n}}{dt} = \kappa(E_{i-1} - E_i)$<br>$\frac{dI}{dt} = \kappa E_n - \delta I$<br>$\frac{dV_{Cell}}{dt} = \pi I - \frac{D(V_{Cell}/I)}{(V_{Cell}/I) + K}V_{Cell}$<br>$\frac{dV_{Sup}}{dt} = \frac{D(V_{Cell}/I)}{(V_{Cell}/I) + K}V_{Cell} - \gamma V_{Sup}$ | | | | | 69.64 |
| | | | | 3 | 53.16<br>53.15<br>53.29<br>53.2<br>53.14<br>53.26 |
| | | | | 5 | 44.48<br>44.33<br>44.02<br>44.83<br>44.85<br>44.44 |
| | | | | 6 | 39.65<br>39.85<br>39.81<br>39.8<br>40.87<br>39.68 |
| | | | | 8 | 33.62<br>33.48<br>32.31<br>33.17<br>33.52<br>33.32 |
| | | | | 10 | 23.12<br>23.17<br>23.05<br>23.16<br>23.16<br>23.09 |
| | | | | 12 | 16.29<br>16.03<br>15.72<br>15.86<br>15.86<br>16.06 |
| | | | | 13 | 13.9<br>13.73<br>13.55<br>13.60<br>13.67<br>13.69 |
| | | | | 14 | 12.69<br>12.77<br>12.59<br>12.64<br>12.65<br>12.53 |

| | | | | | |
|---|---|---|---|---|---|
| | | | | 15 | 12.7<br>12.65<br>12.56<br>12.65<br>12.44<br>12.56 |
| | | | | 16 | 13.22<br>13.21<br>13.16<br>13.15<br>13.11<br>12.97 |
| | | | | 17 | 14.06<br>14.01<br>13.99<br>14.05<br>13.93<br>14.49 |
| | | | | 18 | 15.58<br>15.35<br>15.39<br>16.68<br>15.46<br>15.57 |
| **Density-Dependent Diffusion (per cell)**<br>$\frac{dS}{dt} \cancel{= -\beta_{Sup}V_{Sup}S} - \beta_{Cell}V_{Cell}S$<br>$\frac{dE_1}{dt} = \cancel{\beta_{Sup}V_{Sup}S} + \beta_{Cell}V_{Cell}S - \kappa E_1$<br>$\frac{dE_{i=2,\ldots,n}}{dt} = \kappa(E_{i-1} - E_i)$<br>$\frac{dI}{dt} = \kappa E_n - \delta I$<br>$\frac{dV_{Cell}}{dt} = \pi I - \frac{D(V_{Cell}/I)}{(V_{Cell}/I) + K} V_{Cell}$<br>$\frac{dV_{Sup}}{dt} = \frac{D(V_{Cell}/I)}{(V_{Cell}/I) + K} V_{Cell} - \gamma V_{Sup}$ | $\gamma = 0.04$<br>$\beta_{Sup} = 0$ | $\log_{10}\beta_{Cell}$,<br>$\kappa, \pi, D, K$ | Vero-PCR | 15 | 8.61<br>8.55<br>8.77<br>8.72<br>8.59<br>8.58 |
| **Density-Dependent Diffusion per cell with burst**<br>$\frac{dS}{dt} = -\beta_{Sup}V_{Sup}S - \beta_{Cell}V_{Cell}S$<br>$\frac{dE_1}{dt} = \beta_{Sup}V_{Sup}S + \beta_{Cell}V_{Cell}S - \kappa E_1$<br>$\frac{dE_{i=2,\ldots,18}}{dt} = \kappa(E_{i-1} - E_i)$ | $\gamma = 0.04$ | $\log_{10}\beta_{Sup}$,<br>$\log_{10}\beta_{Cell}$,<br>$f, \kappa, \pi, D, K, \delta$ | Vero-PCR | 1 | 73.69<br>73.99<br>74.16<br>78.95<br>73.87<br>74.32 |
| | | | | 3 | 63.2<br>63.60<br>62.70<br>62.69<br>62.94<br>62.58 |

| | | | | | |
|---|---|---|---|---|---|
| $\frac{dI_1}{dt} = \kappa E_n - \delta I$<br>$\frac{dV_{Cell}}{dt} = \pi I - DV_{Cell}\frac{V_{Cell}}{V_{Cell} + KI} - f\delta I$<br>$\frac{dV_{Sup}}{dt} = DV_{Cell}\frac{V\_Cell}{V_{Cell} + KI} + f\delta I - \gamma V_{Sup}$ | | | | 5 | 47.47<br>46.86<br>56.93<br>58.31<br>46.57<br>47.34 |
| | | | | 6 | 42.5<br>43.00<br>42.75<br>42.62<br>42.83<br>42.91 |
| | | | | 8 | 32.94<br>33.15<br>33.00<br>32.82<br>32.95<br>34.61 |
| | | | | 10 | 22.43<br>22.40<br>22.35<br>22.24<br>22.30<br>22.22 |
| | | | | 12 | 14.44<br>14.31<br>14.20<br>14.49<br>14.35<br>14.27 |
| | | | | 13 | 11.83<br>11.81<br>11.70<br>11.88<br>11.87<br>11.74 |
| | | | | 14 | 10.72<br>10.72<br>10.66<br>10.69<br>10.67<br>10.74 |
| | | | | 15 | 10.66<br>10.56<br>10.56<br>10.70<br>10.48<br>10.56 |

| | | | | | |
|---|---|---|---|---|---|
| | | | | 16 | 11.2<br>11.14<br>11.14<br>11.50<br>11.08<br>11.24 |
| | | | | 17 | 12.17<br>12.13<br>12.14<br>12.26<br>12.16<br>12.55 |
| | | | | 18 | 13.26<br>13.23<br>13.64<br>13.51<br>13.30<br>13.40 |
| **Density-Dependent Diffusion per cell with burst**<br>$\frac{dS}{dt} = \cancel{-\beta_{Sup}V_{Sup}S} - \beta_{Cell}V_{Cell}S$<br>$\frac{dE_1}{dt} = \cancel{\beta_{Sup}V_{Sup}S +} \beta_{Cell}V_{Cell}S - \kappa E_1$<br>$\frac{dE_{i=2,\ldots,18}}{dt} = \kappa(E_{i-1} - E_i)$<br>$\frac{dI_1}{dt} = \kappa E_n - \delta I$<br>$\frac{dV_{Cell}}{dt} = \pi I - DV_{Cell}\frac{V_{Cell}}{V_{Cell} + KI} - f\delta I$<br>$\frac{dV_{Sup}}{dt} = DV_{Cell}\frac{V\_Cell}{V_{Cell} + KI} + f\delta I - \gamma V_{Sup}$ | $\gamma = 0.04$<br>$\beta_{Sup} = 0$ | $\log_{10}\beta_{Cell}$,<br>$f, \kappa, \pi, D, K$ | Vero-PCR | 15 | 6.85<br>6.69<br>7.10<br>6.94<br>6.72<br>**6.65** |
| **Density-Dependent Diffusion (per cell)**<br>$\frac{dE_1}{dt} = -\kappa E_1$<br>$\frac{dE_{i=2,\ldots,n}}{dt} = \kappa(E_{i-1} - E_i)$<br>$\frac{dI}{dt} = \kappa E_n - \delta I$<br>$\frac{dV_{Cell}}{dt} = \pi I - DV_{Cell}\frac{DV_{Cell}/I}{V_{Cell}/I + K}$<br>$\frac{dV_{Sup}}{dt} = DV_{Cell}\frac{DV_{Cell}/I}{V_{Cell}/I + K} - \gamma V_{Sup}$ | $\gamma = 0.04$<br>$\delta = 0.28$ | $\kappa, \pi, D, K$<br>$V_{Cell}(0), V_{Sup}(0)$ | Vero-PFU (High MOI) | 15 | -15.93<br>-58.16<br>-18.35<br>-59.15<br>-19.96<br>-16.34 |

| | | | | | |
|---|---|---|---|---|---|
| **Linear Diffusion**<br>$\frac{dE_1}{dt} = -\kappa E_1$<br>$\frac{dE_{i=2,\dots,n}}{dt} = \kappa(E_{i-1} - E_i)$<br>$\frac{dI}{dt} = \kappa E_n - \delta I$<br>$\frac{dV_{Cell}}{dt} = \pi I - DV_{Cell}$<br>$\frac{dV_{Sup}}{dt} = DV_{Cell} - \gamma V_{Sup}$ | $\gamma = 0.04$<br>$\delta = 0.28$ | $\kappa, \pi, D$<br>$V_{Cell}(0), V_{Sup}(0)$ | Vero-PFU (High MOI) | 15 | -60.77<br>-59.14<br>-60.61<br>-51.26<br>-49.44<br>-15.41 |
| **Linear Diffusion with burst**<br>$\frac{dE_1}{dt} = -\kappa E_1$<br>$\frac{dE_{i=2,\dots,n}}{dt} = \kappa(E_{i-1} - E_i)$<br>$\frac{dI}{dt} = \kappa E_n - \delta I$<br>$\frac{dV_{Cell}}{dt} = \pi I - DV_{Cell} - f\delta I$<br>$\frac{dV_{Sup}}{dt} = DV_{Cell} + f\delta I - \gamma V_{Sup}$ | $\gamma = 0.04$<br>$\delta = 0.28$ | $f, \kappa, \pi, D$<br>$V_{Cell}(0), V_{Sup}(0)$ | Vero-PFU (High MOI) | 15 | -68.52<br>-64.63<br>**-70.82**<br>-60.96<br>-65.33<br>-70.52 |
| **Combined Model with burst**<br>$\frac{dS}{dt} = -\beta_{Cell} V_{Cell,PFU} S$<br>$\frac{dE_1}{dt} = \beta_{Cell} V_{Cell,PFU} S - \kappa E_1$<br>$\frac{dE_{i=2,\dots,n}}{dt} = \kappa(E_{i-1} - E_i)$<br>$\frac{dI}{dt} = \kappa E_n - \delta I$<br>$\frac{dV_{Cell,PFU}}{dt} = \pi_{PFU} I - D_{PFU} V_{Cell,PFU} - f_{PFU}\delta I$<br>$\frac{dV_{Sup,PFU}}{dt} = DV_{Cell,PFU} - \gamma_{PFU} V_{Sup,PFU} + f_{PFU}\delta I$ | $\gamma_I = 0.04$<br>$\gamma_{NI} = 0$<br>$\beta_{Sup} = 0$<br>$\delta = \delta_{PCR}$<br>$\pi_I, \pi_{NI}, D, K, f_{NI}$ | $\log_{10}\beta_{Cell}$,<br>$\kappa, f_I$ | Vero-PCR+PFU | 15 | -20.55<br>-20.60<br>-20.38<br>-20.60<br>-20.42 |

| | | | | | |
|---|---|---|---|---|---|
| $\frac{dV_{Cell,DNA}}{dt} = \pi_{DNA} I - DV_{Cell,DNA} \frac{\frac{V_{Cell,DNA}}{I}}{\frac{V_{Cell,DNA}}{I} + K} - f_{DNA} \delta I$<br>$\frac{dV_{Sup,DNA}}{dt} = DV_{Cell,DNA} \frac{V_{Cell,DNA}/I}{V_{Cell,DNA}/I + K} + f_{DNA} \delta I + \gamma_{DNA} V_{Sup,DNA}$ | | | | | |
| **Combined Model with DNA burst**<br>$\frac{dS}{dt} = -\beta_{Cell} V_{Cell,PFU} S$<br>$\frac{dE_1}{dt} = \beta_{Cell} V_{Cell,PFU} S - \kappa E_1$<br>$\frac{dE_{i=2,\ldots,n}}{dt} = \kappa(E_{i-1} - E_i)$<br>$\frac{dI}{dt} = \kappa E_n - \delta I$<br>$\frac{dV_{Cell,PFU}}{dt} = \pi_{PFU} I - D_{PFU} V_{Cell,PFU}$<br>$\frac{dV_{Sup,PFU}}{dt} = DV_{Cell,PFU} - \gamma_{PFU} V_{Sup,PFU}$<br>$\frac{dV_{Cell,DNA}}{dt} = \pi_{DNA} I - DV_{Cell,DNA} \frac{\frac{V_{Cell,DNA}}{I}}{\frac{V_{Cell,DNA}}{I} + K} - f_{DNA} \delta I$<br>$\frac{dV_{Sup,DNA}}{dt} = DV_{Cell,DNA} \frac{V_{Cell,DNA}/I}{V_{Cell,DNA}/I + K} + f_{DNA} \delta I + \gamma_{DNA} V_{Sup,DNA}$ | $\gamma_I = 0.04$<br>$\gamma_{NI} = 0$<br>$\beta_{Sup} = 0$<br>$\delta = \delta_{PCR}$<br>$\pi_I, \pi_{NI}, D, K, f_{NI}$ | $\log_{10} \beta_{Cell}$, $\kappa$ | Vero-PCR+PFU | 15 | -25.29<br>-25.18<br>-25.29<br>-25.25<br>-25.30<br>-25.34 |

| | | | | | |
|---|---|---|---|---|---|
| **Combined Model with DNA burst with 2 stages of production**<br>$\frac{dS}{dt} = -\beta_{Cell} V_{Cell,PFU} S$<br>$\frac{dE_1}{dt} = \beta_{Cell} V_{Cell,PFU} S - \kappa E_1$<br>$\frac{dE_{i=2,\ldots,n}}{dt} = \kappa(E_{i-1} - E_i)$<br>$\frac{dI_{DNA}}{dt} = \kappa E_n - \lambda I_{DNA}$<br>$\frac{dI_{PFU}}{dt} = \lambda I_{DNA} - \delta I_{PFU}$<br>$\frac{dV_{Cell,PFU}}{dt} = \pi_{PFU} I_{PFU} - D V_{Cell,I}$<br>$\frac{dV_{Sup,PFU}}{dt} = D V_{Cell,PFU} - \gamma_{PFU} V_{Sup,PFU}$<br>$\frac{dV_{Cell,DNA}}{dt}$<br>$= \pi_{DNA}(I_{DNA} + I_{PFU})$<br>$- D V_{Cell,DNA} \frac{\frac{V_{Cell,DNA}}{I_{DNA} + I_{PFU}}}{\frac{V_{Cell,DNA}}{I_{DNA} + I_{PFU}} + K} - f_{DNA} \delta I_{PFU}$<br>$\frac{dV_{Sup,DNA}}{dt} = D V_{Cell,NI} \frac{\frac{V_{Cell,NI}}{I_{DNA} + I_{PFU}}}{\frac{V_{Cell,NI}}{I_{DNA} + I_{PFU}} + K}$<br>$+ f_{DNA} \delta I_{PFU}$<br>$+ \gamma_{PFU} V_{Sup,PFU}$ | $\gamma_I = 0.04$<br>$\gamma_{NI} = 0$<br>$\beta_{Sup} = 0$<br>$\delta = \delta_{PCR}$<br>$\pi_{PFU}, \pi_{DNA}, D_{DN}$,<br>$D_{PFU}\ K, f_{NI}, \kappa$ | $\log_{10} \beta_{Cell}$,<br>$\lambda$ | Vero-PCR+PFU | 15 | **-30.77**<br>-30.98<br>-31.39<br>-29.87<br>-31.02<br>-31.20 |

| | | | | | |
|---|---|---|---|---|---|
| **Combined Model with burst with 2 stages of production**<br>$\frac{dS}{dt} = -\beta_{Cell}V_{Cell,PFU}S$<br>$\frac{dE_1}{dt} = \beta_{Cell}V_{Cell,PFU}S - \kappa E_1$<br>$\frac{dE_{i=2,\dots,n}}{dt} = \kappa(E_{i-1} - E_i)$<br>$\frac{dI_{DNA}}{dt} = \kappa E_n - \lambda I_{DNA} - \delta I_{DNA}$<br>$\frac{dI_{PFU}}{dt} = \lambda I_{DNA} - \delta I_{PFU}$<br>$\frac{dV_{Cell,PFU}}{dt} = \pi_{PFU}I_{PFU} - DV_{Cell,I} - f_{PFU}\delta I_{PFU}$<br>$\frac{dV_{Sup,PFU}}{dt} = DV_{Cell,PFU} - \gamma_{PFU}V_{Sup,PFU} + f_{PFU}\delta I_{PFU}$<br>$\frac{dV_{Cell,DNA}}{dt} = \pi_{DNA}(I_{DNA} + I_{PFU}) - DV_{Cell,NI}\frac{\frac{V_{Cell,NI}}{I_{DNA}+I_{PFU}}}{\frac{V_{Cell,NI}}{I_{DNA}+I_{PFU}} + K} - f_{DNA}\delta(I_{PFU} + I_{DNA})$<br>$\frac{dV_{Sup,DNA}}{dt} = DV_{Cell,NI}\frac{\frac{V_{Cell,NI}}{I_{DNA}+I_{PFU}}}{\frac{V_{Cell,NI}}{I_{DNA}+I_{PFU}} + K} + f_{DNA}\delta(I_{PFU} + I_{DNA}) + \gamma_{PFU}V_{Sup,PFU}$ | $\gamma_I = 0.04$<br>$\gamma_{NI} = 0$<br>$\beta_{Sup} = 0$<br>$\delta = \delta_{PCR}$<br>$\pi_I, \pi_{NI}, D, K, f_{NI}$ | $\log_{10}\beta_{Cell}$, $\lambda, f_{PFU}$ | Vero-PCR+PFU | 15 | -26.74<br>-26.64<br>-26.74<br>-26.84<br>-26.72<br>-26.70 |
| **Combined Model with burst with DNA conversion to PFU**<br>$\frac{dS}{dt} = -\beta_{Cell}V_{Cell,PFU}S$<br>$\frac{dE_1}{dt} = \beta_{Cell}V_{Cell,PFU}S - \kappa E_1$<br>$\frac{dE_{i=2,\dots,n}}{dt} = \kappa(E_{i-1} - E_i)$<br>$\frac{dI_{DNA}}{dt} = \kappa E_n - \delta I_{DNA}$ | $\gamma_I = 0.04$<br>$\gamma_{NI} = 0$<br>$\beta_{Sup} = 0$<br>$\delta = \delta_{PCR}$<br>$\pi_{DNA}, D_{DNA}, D_{P}$[illegible] | $\log_{10}\beta_{Cell}$, $\alpha$ | Vero-PCR+PFU | 15 | -26.00<br>-26.05<br>-25.9<br>-25.96 |

| | | | | | |
|---|---|---|---|---|---|
| $\frac{dV_{Cell,DNA}}{dt} = \pi_{DNA}(I_{DNA}) - DV_{Cell,NI}\frac{\frac{V_{Cell,NI}}{I_{DNA}}}{\frac{V_{Cell,NI}}{I_{DNA}}+K} - f_{DNA}\delta I_{DNA}$<br>$\frac{dV_{Sup,DNA}}{dt} = DV_{Cell,NI}\frac{\frac{V_{Cell,NI}}{I_{DNA}}}{\frac{V_{Cell,NI}}{I_{DNA}}+K} + f_{DNA}\delta I_{DNA} + \gamma_{PFU}V_{Sup,PFU}$<br>$\frac{dV_{Cell,PFU}}{dt} = \alpha V_{Cell,DNA} - DV_{Cell,PFU}$<br>$\frac{dV_{Sup,PFU}}{dt} = DV_{Cell,PFU} - \gamma_{PFU}V_{Sup,PFU}$ | | | | | |
| **Linear Diffusion**<br>$\frac{dS}{dt} = -\beta_{Sup}V_{Sup}S - \beta_{Cell}V_{Cell}S$<br>$\frac{dE_1}{dt} = \beta_{Sup}V_{Sup}S + \beta_{Cell}V_{Cell}S - \kappa E_1$<br>$\frac{dE_{i=2,\ldots,n}}{dt} = \kappa(E_{i-1} - E_i)$<br>$\frac{dI}{dt} = \kappa E_n - \delta I$<br>$\frac{dV_{Cell}}{dt} = \pi I - DV_{Cell}$<br>$\frac{dV_{Sup}}{dt} = DV_{Cell} - \gamma V_{Sup}$ | $\gamma = 0.04$<br>$\delta = 0$ | $\log_{10}\beta_{Cell}$, $\log_{1}$[illegible]<br>$\kappa, \pi, D$, $S_0$ | N2A-PCR | 1 | 52.01<br>52.06<br>52.22<br>51.86<br>52.18<br>52.18 |
| **Linear Diffusion**<br>$\frac{dS}{dt} = -\beta_{Sup}V_{Sup}S - \beta_{Cell}V_{Cell}S$<br>$\frac{dE_1}{dt} = \beta_{Sup}V_{Sup}S + \beta_{Cell}V_{Cell}S - \kappa E_1$<br>$\frac{dE_{i=2,\ldots,n}}{dt} = \kappa(E_{i-1} - E_i)$ | $\gamma = 0.04$<br>$\delta = 0$ | $\log_{10}\beta_{Cell}$, $\log_{1}$[illegible]<br>$\kappa, \pi, D$, $S_0$<br>$\log_{10}\beta_{Cell}$<br>$\kappa, \pi, D$, $S_0$ | N2A-PCR | 3 | 74.99<br>55.27<br>75.06<br>55.49<br>54.71<br>55.35 |
| | $\gamma = 0.04$<br>$\delta = 0$ | | | 5 | 55.84<br>55.90<br>55.98 |

| | | | | | |
|---|---|---|---|---|---|
| $\frac{dI}{dt} = \kappa E_n - \delta I$<br>$\frac{dV_{Cell}}{dt} = \pi I - DV_{Cell}$<br>$\frac{dV_{Sup}}{dt} = DV_{Cell} - \gamma V_{Sup}$ | | | | | 56.69<br>55.87<br>56.03 |
| | | | | 7 | 56.49<br>56.59<br>56.64<br>56.45<br>56.56<br>56.52 |
| **Linear Diffusion**<br>$\frac{dS}{dt} = \cancel{-\beta_{Sup}V_{Sup}S} - \beta_{Cell}V_{Cell}S$<br>$\frac{dE_1}{dt} = \cancel{\beta_{Sup}V_{Sup}S} + \beta_{Cell}V_{Cell}S - \kappa E_1$<br>$\frac{dE_{i=2,\ldots,n}}{dt} = \kappa(E_{i-1} - E_i)$<br>$\frac{dI}{dt} = \kappa E_n - \delta I$<br>$\frac{dV_{Cell}}{dt} = \pi I - DV_{Cell}$<br>$\frac{dV_{Sup}}{dt} = DV_{Cell} - \gamma V_{Sup}$ | | | | 1 | **47.89**<br>47.93<br>48.13<br>48.25<br>48.31<br>48.35 |
| **Linear Diffusion**<br>$\frac{dS}{dt} = \cancel{-\beta_{Sup}V_{Sup}S} - \beta_{Cell}V_{Cell}S$<br>$\frac{dE_1}{dt} = \cancel{\beta_{Sup}V_{Sup}S} + \beta_{Cell}V_{Cell}S - \kappa E_1$<br>$\frac{dE_{i=2,\ldots,n}}{dt} = \kappa(E_{i-1} - E_i)$<br>$\frac{dI}{dt} = \kappa E_n - \delta I$<br>$\frac{dV_{Cell}}{dt} = \pi I - DV_{Cell}$<br>$\frac{dV_{Sup}}{dt} = DV_{Cell} - \gamma V_{Sup}$ | $\gamma = 0.04$<br>$\delta_I = 0$ | $\log_{10}\beta_{Cell}$<br>$\kappa, \pi, D$ | N2A-PCR | 1 | 49.40<br>49.53<br>49.47<br>49.35<br>49.40<br>49.49 |

| | | | | | |
|---|---|---|---|---|---|
| **Density-dependent Diffusion**<br>$\frac{dS}{dt} = \cancel{-\beta_{Sup}V_{Sup}S} - \beta_{Cell}V_{Cell}S$<br>$\frac{dE_1}{dt} = \cancel{\beta_{Sup}V_{Sup}S} + \beta_{Cell}V_{Cell}S - \kappa E_1$<br>$\frac{dE_{i=2,\dots,n}}{dt} = \kappa(E_{i-1} - E_i)$<br>$\frac{dI}{dt} = \kappa E_n - \delta I$<br>$\frac{dV_{Cell}}{dt} = \pi I - DV_{Cell}\frac{V_{Cell}}{V_{Cell} + KI}$<br>$\frac{dV_{Sup}}{dt} = DV_{Cell}\frac{V_{Cell}}{V_{Cell} + KI} - \gamma V_{Sup}$ | $\gamma = 0.04$<br>$\delta_I = 0$ | $\log_{10}\beta_{Cell}$<br>$\kappa, \pi, D, K, S_0$ | N2A-PCR | 1 | 51.89<br>52.22<br>52.38<br>52.68<br>52.26<br>52.42 |

*all runs converged to the same global minima with the same AIC.

**Supplementary Table S2. List of competing models with the fixed and estimated parameters**. Each model was run six times with different initial guesses for the parameters and the AIC score of each run is recorded here.

| parameter | value | | Estimated at | Transferred to stage 3? |
|---|---|---|---|---|
| | MOI=3 | MOI=0.01 | | |
| $\log_{10}\beta_{Cell}$ | -7.22 | -7.22 | Stage 1 | No |
| $\pi_{DNA}$ | 5250.96 | 5252.02 | Stage 1 | Yes |
| $D_{DNA}$ | 0.047 | 0.047 | Stage 1 | Yes |
| $\kappa_{DNA}$ | 1.37 | 0.82 | Stage 1 | Yes |
| $\log_{10}K$ | 6.37 | 6.38 | Stage 1 | Yes |
| $\delta$ | 0.28 | 0.072 | Stage 1 | Yes |
| $f_{DNA}$ | 155.92 | 65.08 | Stage 1 | Yes |
| $\pi_{PFU}$ | 23.47 | N/A | Stage 2 | Yes |
| $D_{PFU}$ | 0.003 | N/A | Stage 2 | Yes |
| $f_{PFU}$ | 0.3(=0) | N/A | Stage 2 | Yes |
| $\kappa_{PFU}$ | 0.97 | N/A | Stage 2 | No |
| $\log_{10}\beta_{Cell,PFU}$ | -4.38 | -4.33 | Stage 3 | estimated |
| $\lambda$ | 0.53 | 0.53 | Stage 3 | estimated |
| $\gamma_{PFU}$ | 0.04 | 0.04 | fixed | Yes |
| $\gamma_{DNA}$ | 0 | 0 | fixed | Yes |

**Supplementary Table S3. Parameter values estimated at each stage of modeling Vero cell data.**

| id | $\pi$ | D | $\gamma$ | $\log_{10}\beta_{Cell}$ | $\kappa$ | $S_0$ |
|---|---|---|---|---|---|---|
| High | 1027.59 | 0.0083 | 0.04 | -11.01 | 0.024 | 11790.47 |
| Low | 1026.96 | 0.0083 | 0.04 | -11.01 | 0.024 | 974274.69 |

**Supplementary Table S4. Parameter values are estimated by fitting the model to N2A PCR data.** All parameters are consistent between the high and low MOI experiments. The initial number of susceptible cells is estimated with MOI as the covariate for $S_0$. Dark grey cells mark the fixed parameters. Highlighted parameters were unidentifiable.

| quantity | Value(units) | method |
|---|---|---|
| **VERO Cell model stage 3 (final model)** | | |
| Average time spent in eclipse phase | 10.9 hrs (high MOI)<br>18.29 hrs (low MOI) | $n/\kappa$ |
| How much time ~90% of cells spend in the eclipse phase | 7-16 hrs (high MOI)<br>11-27 hrs (low MOI) | 90% confidence interval equal areas around median |
| Earliest time that viral DNA production begins | 2.5hrs (high MOI)<br>6.5 (hrs) (low MOI) | The first t at which $I_{DNA}(t)>=1$ |
| Earliest time that infectious viral particles production begins | 3 hrs (high MOI)<br>7.2 (hrs) (low MOI) | The first t at which $I_{PFU}(t)>=1$ |
| %90 of cells entering productive phase by | ~15 hrs (high MOI)<br>~24.5 hrs (low MOI) | Gamma.cdf(t)~0.9 |
| time of egress of first infectious particles **post infection** | 8.0 hrs (high MOI)<br>20.5 hrs(low MOI) | The first t at which $V_{Sup,I}(t)$ is at least 1 copy/ml larger than $V_{Sup,I}(t)$ when DNA production starts (when $I_{DNA}(t)>=1$) |
| time of egress of first Infectious particles **post DNA production** | 8-2.5=5.5(hrs)high MOI<br>20.5-6.5= 14 (hrs)low MOI | |
| time of egress of first viral DNA (non-infectious model compartment) particles **post infection** | 2.5 hours (high MOI)<br>6.5 hours (low MOI) | The first t at which $V_{Sup,DNA}(t)$ is at least 1 copy/ml larger than $V_{Sup,DNA}(t)$ when DNA production starts (when $I_{DNA}(t)>=1$) |
| time of egress of first viral DNA non-infectious (model compartment)particles **post production** | immediately | |
| Number of HSV DNA produces for each infectious particle | 5250.96/23.47 = 223.7 | $\pi_{DNA}/\pi_{PFU}$ |
| Half-life of productively infected cells (after start of production) | ~3.5 (hrs) (high MOI)<br>~14 (hrs) (low MOI) | $1/\delta$ |
| Half-life of productively infected cells (after start of infection) | ~14.4 (hrs) (high MOI)<br>~32.29(hrs) (low MOI) | $\frac{n}{\kappa}+\frac{1}{\delta}$ |
| In low MOI 95% of cells become infected (leave susceptible cells) by ….. | ~16.25 (hrs) | t at which the susceptible cell population is less than equal $0.05*3x10^5$ |
| proportion of cells that entered the productive stage early (before 18 hours) | 0.15% (low MOI) | I(t=18)/30000 |
| **N2A Cell Model vs Vero Cell model stage 1 (PCR model)** | | |
| Average time spent in eclipse phase (Vero cells) | 14.15 (hrs) | $\frac{n}{\kappa_{Pop}}$ |
| Variability of time spent in eclipse phase (Vero cells) | 3.65 hrs | Standard deviation of the gamma(n, $\kappa$) = $\sqrt{\frac{n}{\kappa_{pop}^2}}$ |
| Average time spent in eclipse phase (N2A cells) | 41.6 ~ 42 (hrs) | $\frac{n}{\kappa}$ |
| Variability of time spent in eclipse phase (Vero cells) | 41.6 hrs | Standard deviation of the gamma(n, $\kappa$) = $\sqrt{\frac{n}{\kappa^2}}$ |

**Supplementary Table S4**. Measured quantities using model parameters.
